# Inferring Cell Fate Trajectories in Time-Resolved Metabolic RNA Labeling data

**DOI:** 10.64898/2026.06.11.731638

**Authors:** Anna Audit, Gabriel Peyré, Laura Cantini

## Abstract

Single-cell RNA sequencing provides high-resolution snapshots of cellular states but lacks direct information about transcriptional dynamics. Metabolic RNA labeling addresses this limitation by distinguishing newly synthesized RNA, offering insight into the direction of cell state changes, and providing valuable information when attempting to recover the underlying continuous dynamics from static snapshots of cell distributions. However, existing trajectory inference methods do not fully exploit this additional signal. Here, we propose FLOWSATATE, a framework for single-cell trajectory inference that leverages time-resolved RNA labeling within an Optimal Transport setting. We model cell dynamics as a gradient flow in an inferred potential landscape parameterized by a neural network, integrating both total and labeled RNA across time points. The learned potential enables identification of key genes and transcription factors driving cell fate decisions and supports prediction of future cellular states. We benchmark our approach on its ability to generalize unseen data and recover coherent trajectories. We also apply it to study colorectal cancer response to demethylation treatment as well as neuronal differentiation of embryonic stem cells.

## Main

Single-cell RNA sequencing technology has revolutionized the biomedical field by enabling the high-resolution analysis of the transcriptional content of individual cells^1^. Despite its significant advancements, single-cell RNA sequencing (scRNA) provides only a “static” snapshot of cellular states, lacking the ability to capture transcriptional dynamics. Metabolic RNA labeling technologies address this limitation by enabling simultaneous measurement of total RNA and newly synthesized RNA within each cell. These approaches typically rely on the incorporation of 4-thiouridine (4sU) into nascent transcripts, allowing the distinction between newly transcribed and pre-existing RNA ^2–4^. The labeling of newly synthesized RNA provides simultaneous access to the cell’s gene expression (“total RNA”) as well as local insights into the direction of evolution of each individual cell (“labeled RNA”), which traditional sequencing techniques cannot capture. When metabolic RNA labeling sequencing is done at multiple time points we additionally have access to the evolution of the entire cell population through the successive snapshots. However, this information is unpaired as sequencing is destructive. This addition of simultaneous information on the level of total and labeled RNA at the cell level coupled with snapshots of the evolution of the full population adds key information for inferring cellular trajectories.

Trajectory inference methods aim to recover continuous dynamics from static snapshots of the cell distribution ^5^. A broad class of methods has been developed to address this challenge ^6^. When only one time point is available, some methods aim to situate cells along a trajectory based on a pseudotime inferred from their transcriptomic profiles^7–9^. For instance, Monocle^9^ computes cell pseudotime as the distance along a graph learned from transcriptomic profiles. However those methods infer the cell’s position on a purely computational axis which does not always reflect actual temporal ordering^1^. Recently, the concept of RNA velocity has been proposed ^10–13^. RNA velocity methods estimate the local directionality of a cell’s evolution by comparing the abundance of unspliced (nascent) transcripts and spliced (mature) transcripts for each gene. Velocity methods rely on simplified kinetic models and may also struggle to capture the intricate dynamics of individual cells ^1^. To overcome the limitations of single-time point data, methods leveraging multiple time points have been proposed. In particular, approaches based on Optimal Transport (OT) model the evolution of cell populations as transport between distributions across time^14,15,16^. In particular, Wasserstein gradient flows, first introduced by Jordan, Kinderlehrer, and Otto through their connection with the Fokker–Planck equation, have been recently applied to trajectory inference in single-cell transcriptomics ^17–21^. These methods are trained on recovering the correct full cell population provided by scRNA at different time points. As RNA labeling has only recently become available, current methods do not incorporate the additional information that it provides. Information derived from labeled RNA can instead be included in the training objective as a cell-level prior on the direction of a cellular evolution, in addition to the population-level constraints provided by the observed cell distributions.

We developed FLOWSTATE, a method for single-trajectory inference that leverages information from time-resolved RNA labeling. FLOWSTATE introduces methodological developments that explicitly integrate labeled RNA measurements in its objective while allowing reconstruction of future transcriptional states and improving the inference of cellular dynamics across time-resolved snapshots. FLOWSTATE relies on the Optimal Transport formalism, which has increasingly been used in single-cell genomics as it enables meaningful comparisons between unpaired probability distributions through Wasserstein distances. The Wasserstein gradient flow is a commonly used framework to model the evolution of a distribution in an energy landscape^17,19^. As has been previously done we adopt a framework based on the assumption that cellular dynamics are governed by an underlying potential function and learn this driving potential^18,20^. Modeling cellular differentiation as the minimization of such a potential originates from systems biology and Waddington’s epigenetic landscape concept ^22^. Understanding this potential can reveal the genes and transcription factors driving cellular evolution, uncover multiple possible trajectories, and predict a cell’s future state from its initial condition.

We benchmarked FLOWSTATE on its ability to predict unseen transcriptional states as well as recover coherent cell type transitions against state-of-the-art methods that do not use labeled RNA information. When applied to HCT116 colorectal cells treated with a DNA demethylating agent ^4^, FLOWSTATE recovered known effects of the demethylating treatment as well as candidate genes of potential therapeutic relevance, such as *FLI1*. We also applied FLOWSTATE to a dataset of neuronal differentiation of mouse embryonic stem cells ^3^. Our results suggest that the *Slit/Robo* signaling axis is established during motor neurons commitment, earlier than previously appreciated.

## Results

### 1. FLOWSTATE: a new tool for trajectory inference from time-resolved metabolic RNA labeling

FLOWSTATE learns a potential informed by both the single cell RNA snapshots as well as the dynamic information provided by the labeled RNA. This potential can then be used to predict the evolution of the cells and give insight into the genes and transcription factors driving cellular dynamics.

FLOWSTATE takes as input cell populations profiled at multiple time points together with their total and labeled RNA measurements. (Figure 1A). At every time point, we thus have two count matrices available. The total RNA is the sum of old and new RNA, corresponding to the count matrix that standard single-cell RNA sequencing would provide. The second count matrix, containing exactly the same set of cells, corresponds to the counts of newly transcribed RNA that have been labeled (i.e. labeled RNA).

**Figure 1.**
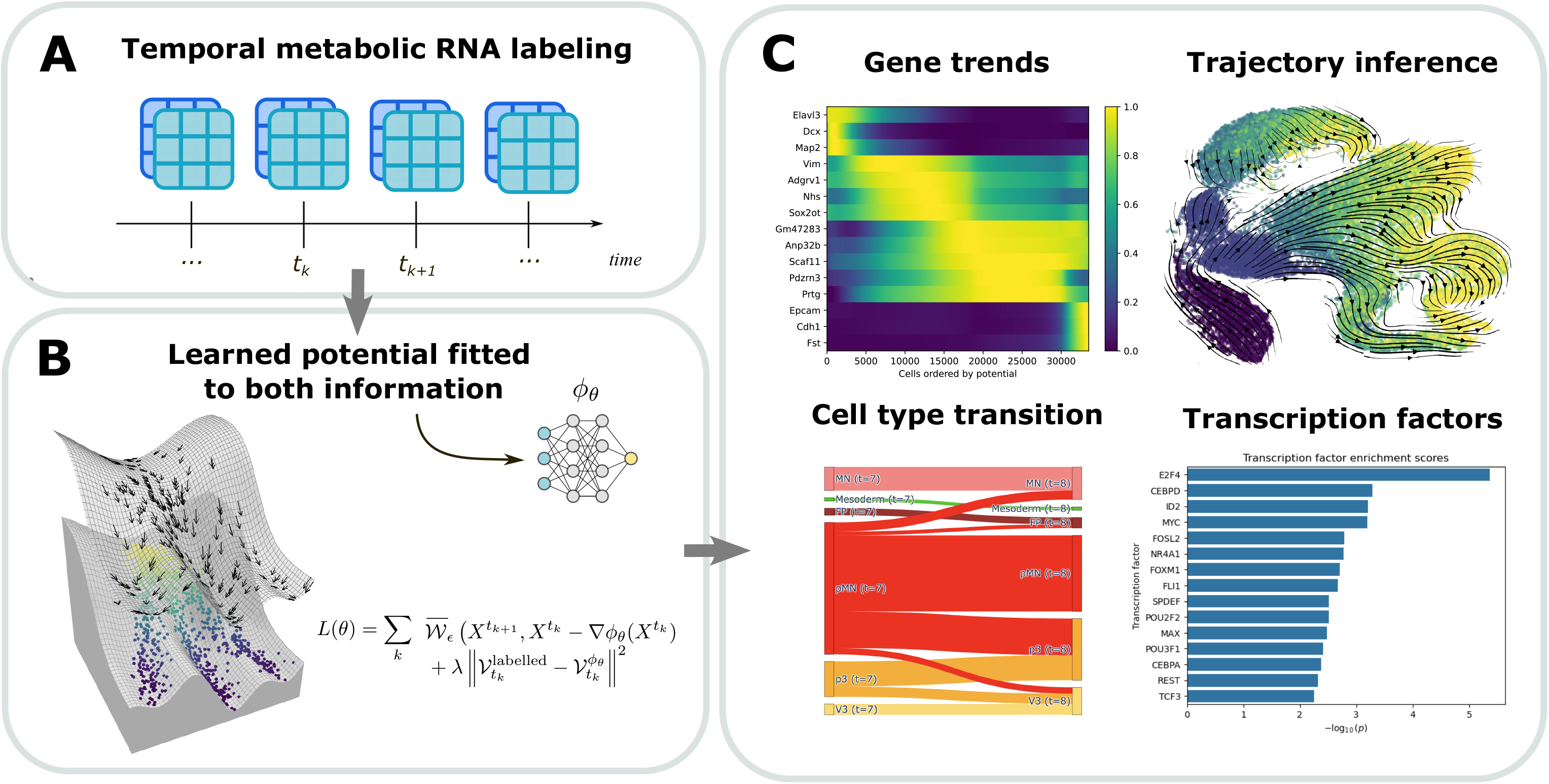
The FLOWSTATE workflow. A. FLOWSTATE takes as input cell populations profiled at multiple time points together with their total and labeled RNA measurements. B. FLOWSTATE learns a potential informed by both the total and labeled RNA C. After training, the potential can be used to reveal key genes and transcription factors, perform trajectory inference, and obtain cell type transitions.

The method learns the parameters *θ* of a neural network *ϕ*_*θ*_ that assigns a potential value to each cell based on its gene expression profile *x* (Figure 1B). This potential defines a Waddington-like landscape^22^ in which undifferentiated or early states lie in regions of high potential and cells evolve toward lower-potential transcriptional states corresponding to more mature states. The potential is estimated by minimizing a joint objective *L*(*θ*) combining a regularized Wasserstein loss with an euclidean discrepancy term. The Wasserstein loss compares the predicted and observed full cell populations while the euclidean term compares the dynamic predicted by the model with the one observed from labeled RNA.

The newly synthesized labeled RNA only captures transcriptional production. However the cell’s evolution is the combined effects of RNA production and degradation, for this reason we need to correct the labeled RNA to account for gene degradation. The experimental protocol varies across datasets. In some settings^23,24^, the beginning of the labeling coincides with the sequencing time of a previous snapshot. In this case, we design an approach that exploits labelled RNA to reconstruct each cell’s earlier transcriptional state, specifically its state at the beginning of labeling. These degradation rates are estimated using an optimal transport objective that minimizes the discrepancy between the estimated cell distribution at the beginning of labeling and the observed ground-truth cell distribution at the corresponding earlier time point (see Online Methods and Supp Figure 1). In other settings^3,4^, the labeling duration (e.g., 2h) is shorter than the time interval between snapshots and no matching exists between the beginning of the labeling and earlier time points. In this case, we will consider that the newly synthesized RNA, once corrected for degradation rates, provides an instantaneous estimate of the cell’s transcriptional velocity. After this correction is done, the scaled, normalized, and log-transformed expression matrix is projected into PCA space to make the framework computationally tractable.

We have access to the cell distribution for a collection of time points *t*_1_ < … < *t*_*k*_ < … <*t*_*K*_. For any time point *t*, we denote by 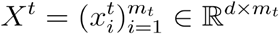 the corresponding dataset, where *m*_*t*_ is the number of cells and *d* is the dimension of the representation space. The vector 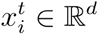 represents the gene expression profile of the *i*-th cell belonging to time *t*. We also observe labeled RNA, denoted by *n*. This corresponds to RNA transcripts produced during a labelling interval of duration *τ* before sequencing. Similarly, we denote 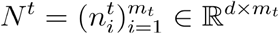 where the vector *n*_*i*_ ∈ ℝ^*d*^ represents the labeled RNA of the *i*-th cell. We define the empirical distribution associated with time point *t* as 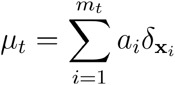, a discrete distribution *m*_*t*_ of cells with weights *a*_*i*_ satisfying 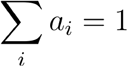.The *a*_*i*_ weights can either be chosen to be uniform and equal to 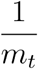 or be set to a computed growth rate for each cell. This has been proposed in^14,16^ to take into account the fact that cells with larger growth rate should have more descendents.

We train the neural network *ϕ*_*θ*_ by predicting the distribution of cells at time point *t*_*k*+1_ from the distribution 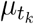 at time *t*_*k*_, comparing this prediction to the ground truth using our joint objective *L*(*θ*)and updating the network parameters accordingly. We denote by 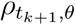 the distribution at time *t*_*k*+1_ predicted from 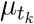 by FLOWSTATE with parameters *θ*. In our case 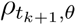 is obtained by moving each cell independently along the gradient of the potential so that 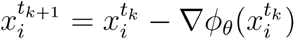. Existing approaches^17,18^ using potentials compare predicted and observed distributions using the Sinkhorn divergence^25^ 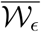 (see online methods); however, this formulation does not incorporate information from newly synthesized RNA.

The new loss we propose is the following :

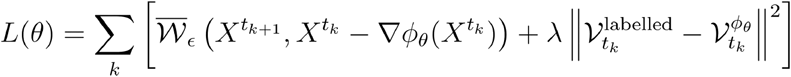

The first term of the loss 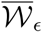 corresponds to the Sinkhorn divergence previously used in ^17,18,20^ .The second *ℓ*^2^ term aligns the dynamics predicted by the learned potential with dynamical information derived from labeled RNA measurements. Here, 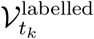 represents an estimate of the cellular evolution inferred from labeled RNA, while 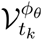 denotes the evolution predicted by the learned Wasserstein gradient flow. Depending on the experimental setting, these quantities may correspond either to instantaneous velocities or to accumulated trajectories across multiple time steps. Unlike the optimal transport term (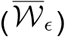), which compares entire population distributions in an unpaired manner, this supervision term leverages paired cell-level dynamical information obtained from labeled RNA measurements. The discretization scheme, explicit integration and neural architecture follows previous implementation done in STORIES^20^ . The parameter *λ* controls the relative importance assigned to this paired dynamical supervision with respect to the optimal transport objective. *λ* = 0 corresponds to a model that does not incorporate labeled RNA information and serves as our baseline model to evaluate the added value of the information provided by labeled RNA.

Once training is complete several interpretable outputs can be derived from the potential (Figure 1C). The gradient of the potential can be used as a cell’s velocity. Gene and transcription factors driving the evolution can be identified as those most correlated with the potential and cell type transitions across time points can also be obtained.

FLOWSTATE is implemented as an open-source Python package integrated into the scverse ^26^ ecosystem and provides visualization tools to explore gene trends, transcription factor dynamics, and cell-type transition probabilities derived from the learned potential. The code is available at https://github.com/cantinilab/FLOWSTATE.

In the next section, we present benchmarking results, where performance is evaluated on the prediction of previously unseen time points and compared against PRESCIENT^18^, the only state-of-the-art method directly comparable on our dataset.

### 2. FLOWSTATE overperforms methods that do not incorporate RNA labeling information

We benchmark FLOWSTATE on three time-resolved metabolic RNA labeling data (Figure 2A). The first dataset studies the response of lung cancer cells to treatment. Lung adenocarcinoma-derived A549 cells were treated with dexamethasone (DEX) and sequenced after 0, 2, 4, 6, 8, and 10h of treatment with a labeling duration of 2h at each time point ^23^. In the this dataset, because the labeling period matches the interval between time points (2h), for any time *t* we can reconstruct the state of each cell at the immediately preceding time point. The second dataset follows the temporal evolution of excitatory neurons. Primary mouse cortical cultures were stimulated for different durations of neuronal activity (0, 15, 30, 60, and 120 min), again following a 2h labeling period ^24^. Since the population is assumed to remain stable prior to stimulation, we consider the distribution at the beginning of labeling for each time point to be equivalent to the unstimulated distribution (0 min). Thus for any time *t* we reconstruct the state of every cell at the very first time point (0min). These two datasets correspond to the setting in which we can reconstruct each cell’s transcriptional state at a previous time point. The third dataset studies the development of mouse embryonic stem (ES) cells differentiating into neural tube identities; cells were sequenced daily from day 3 to day 8 following a 2h labeling period ^3^. This corresponds to the setting where the labeling duration is much shorter than the interval between time points, and the labeled RNA is used to estimate an instantaneous velocity.

**Figure 2.**
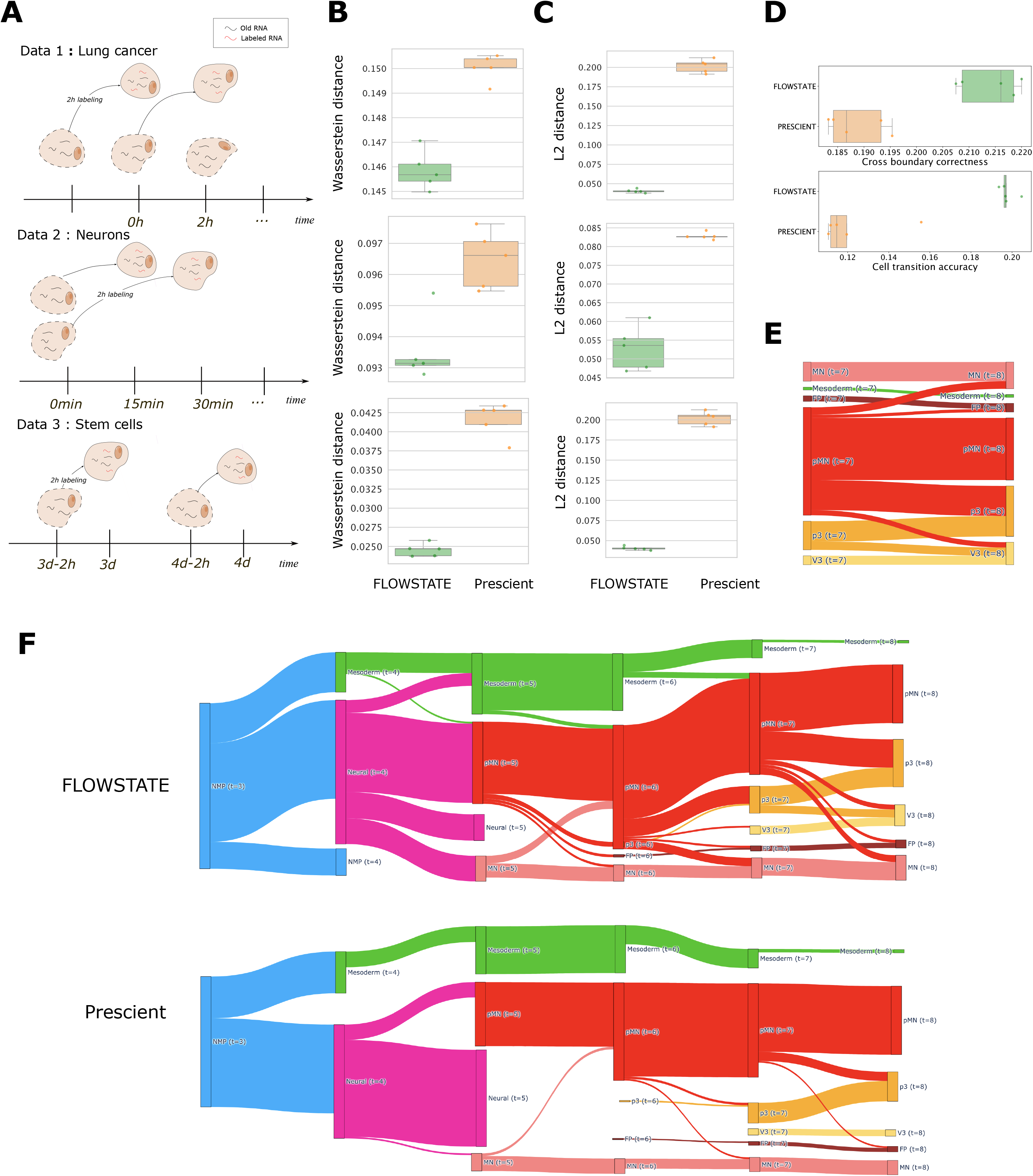
Benchmark of FLOWSTATE. A. Visual representation of the experimental setup of the three datasets used in the benchmarking B. Wasserstein loss between prediction and ground truth at the last time point. Scores are reported for n=5 seeds. C. L2 loss between prediction and ground truth at the last time point. Scores are reported for n=5 seeds D. Cell type transition scores. Scores are reported for n=5 seeds E. Visualization of cell type transitions at the last time point F. Visualization of cell type transitions across time for FLOWSTATE and PRESCIENT.

We evaluate FLOWSTATE on its ability to predict the gene expression in unseen time points. During training, the last time point is removed from the training set. We then test the model’s ability to generate accurate predictions for the last unseen data. This is particularly challenging because it contains cell profiles never seen during training. For the embryonic stem cell data set where three replicates are available we also evaluate FLOWSTATE on the prediction quality for full second and third replicates after training only on the first.

The predicted distribution is compared to the ground truth using the 2-Wasserstein distance (see online methods) on total RNA and the *ℓ*^2^ loss on labeled RNA. The Wasserstein loss assesses whether the overall target distribution is correctly recovered from the predicted distribution. The *ℓ*^2^ loss measures how well the inferred potential aligns with the velocity derived from labeled RNA. For the embryonic stem cell dataset, where multiple cell types are present, we also evaluate the accuracy of the cell type transitions derived from the potential with respect to known transitions across all time points. This provides insights into whether the inferred individual evolutions are coherent with the biological knowledge. In the absence of ground truth cell pairing in between the time point this provides the best accessible ground truth. In order to obtain transitions between cell types from one time point to the next, each cell is first pushed forward in time by the model, which gives a predicted future position. We then classify the predicted future state using a k-nn classifier trained on all the data cell types. By aggregating over all the cells from one time point, this gives us the transition probability across time points (see online methods). Note that when predicting the transition from *t*_*k*_ to *t*_*k*+1_ we do not force the cell type distribution at time to *t*_*k*+1_ match with ground truth cell type distribution at this time point. This method allows for final cell types such as mesoderm to only be paired with themselves even when the abundance of that cell type varies across time. The cell-type transitions are compared with the known transitions using two scores: the cell type transition accuracy between time points corresponding to the proportion of correct transitions and the Cross-Boundary Directionality (CBD), evaluating whether the velocity field derived from the potential points is coherent across cell-type boundaries.

To evaluate the contribution of labeled RNA information, we benchmark FLOWSTATE for multiple values of the *λ* parameter between 0 and 1 and assess the performances against the baseline model that does not take labeled RNA (meaning *λ =* 0). Across all three datasets, the same trend is observed : introducing the *ℓ*^2^ coupling term initially improves the prediction of unseen snapshots under the Wasserstein and Sinkhorn metrics (see Supp Figure 2 and Supp Figure 3) whereas excessively large values degrade performance. This shows the existence of a tradeoff between the information provided by the total and labeled RNA. The value *λ =* 0.25 consistently performs better than the baseline in all datasets and is either the best or second-best choice in every case (see Supp Figure 2). We therefore use *λ =* 0.25 in subsequent analyses.

With this optimal value, FLOWSTATE predicts the final unseen time point more accurately than PRESCIENT across all datasets (Figure 2B) and also systematically performs better on all time points replicates (see Supp Figure 4) . The *ℓ*^2^ loss is also always lower for FLOWSTATE (Figure 2C), this task is easier to achieve for FLOWSTATE as it explicitly optimizes on this term. We further evaluate cell-type transition accuracy on the stem cell dataset, where FLOWSTATE outperforms PRESCIENT both in transition accuracy and CBD metric (Figure 2D). Finally, also looking qualitatively at the cell type transitions (Figure 2E and F), those inferred by FLOWSTATE are more biologically coherent, showcasing the transition from neuro-mesodermal progenitors (NMPs) cells to neural and mesoderm cell type and then the transition from neural to three possible fates: motorneurons (MN), floorplate (FP), or p3 cells. PRESCIENT, on the other hand, misses some key processes like the emergence of the pMN, P3 and FP populations.

### 3. FLOWSTATE highlights the downstream effects of 5-aza-CdR and identifies potential treatment targets in colorectal cancer HCT116 cells

We apply FLOWSTATE to HCT116 colorectal cancer cells treated with 5-aza-2′-deoxycytidine (5-aza-CdR), a DNA demethylating agent. 5-aza-CdR has demonstrated clinical efficacy as an anti-tumor therapy, although the mechanisms underlying its tumor-suppressive effects remain incompletely understood. Its induction of global DNA demethylation can reactivate previously silenced tumor suppressor genes, while also promoting cellular senescence through activation of cell cycle arrest^27,28^. However, the therapeutic effect of 5-aza-CdR could also be offset by reactivating previously silenced oncogenes (^29,30^). The evolution of colorectal cancer cells in early stages (first three days) of treatment by 5-aza-CdR remains largely unknown due to the lack of methods that capture dynamics over a short period of time. In particular, understanding whether the global demethylation also leads to the reactivation of previously silenced oncogenes and genes that promote proliferation and could thus offset the therapeutic effect of 5-aza-CdR. To better understand this process we consider a dataset composed of HCT116 cells treated at a low dose (300 nM) of 5-aza-CdR for varying duration 0, 1, 2, and 3 days ^4^.

We use FLOWSTATE to reconstruct the cellular evolution induced by 5-aza-CdR. Day 3 cells show an expression profile in between the one of cells in day 1 and day 2. The velocity field derived from the potential recovers an equilibrium state at day 2 toward which the cells from day 0, day 1 and day 3 move, as in the original publication ^4^ (Figure 3A).

**Figure 3.**
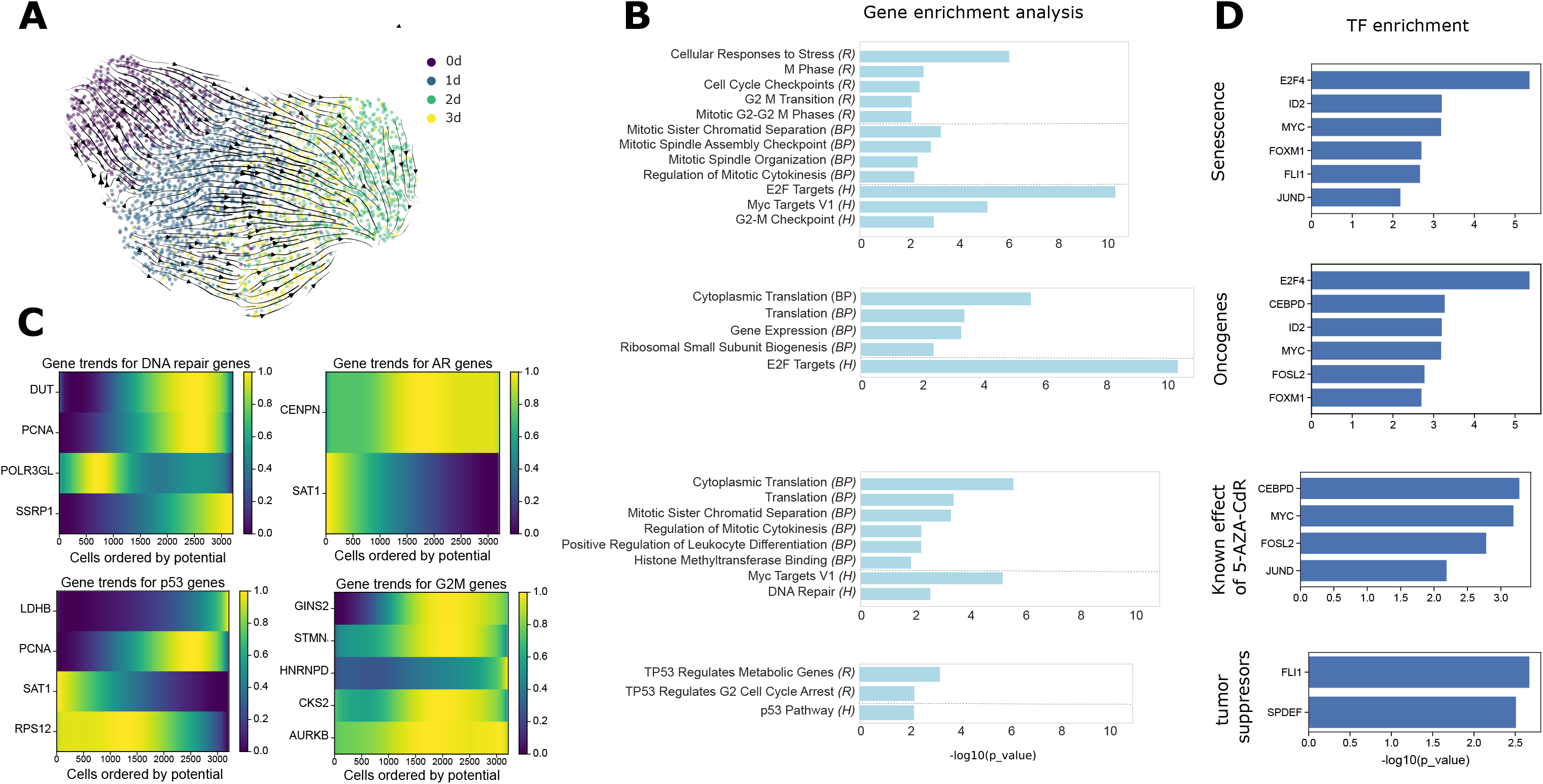
Trajectory inference on colorectal cancer cells. A. Velocity derived from the potential computed by FLOWSTATE plotted in Umap B. Gene enrichment analysis (*R* : Reactome, *BP* : GO Biological Process, *H*: MSigDB Hallmark) performed on genes most correlated with the potential computed by FLOWSTATE. Expression regressed using a spline model along the potential. C. Normalized gene expression for the genes most correlated with the potential computed by FLOWSTATE. Expression regressed using a spline model along the potential. Cells are ordered with potential decreasing from right to left, putting more mature cells on the left. D. Enrichment score of relevant transcription factors targeting the genes most correlated with the potential. A one-sided Wilcoxon rank-sum test is used to report P values.

We then explore the transcription factors and genes driving the evolution of the cells. The potential inferred by our method provides an interpretable measure of cellular differentiation, associating a lower potential with more differentiated cells. To identify genes driving these dynamics, we regress each gene’s expression against the potential using a spline regression model ^31^, and define high-scoring genes as candidate drivers (see online methods). We select our driver genes as those that have the highest correlation with the potential while also selecting genes whose peak of expression is located at different points of the trajectory (beginning, middle, or end) to ensure that we capture the full dynamics. Performing enrichment on these genes (see online methods) shows them to be associated mainly with cell cycle regulation and senescence, as well as oncogenes and tumor suppressors (Supp table 3). Among the top enriched processes, we identify pathways associated with the regulation of the cell cycle (Figure 3B). In particular, we identify G2/M checkpoint, a control point in the cell cycle that ensures cells only enter mitosis after any DNA damage has been properly repaired. Genes associated with the G2/M checkpoint are mostly activated at the beginning of the treatment as a response to global demethylation (Figure 3C). We also find that several processes linked to the tumor suppressor TP53 (p53) are also significantly enriched (Figure 3B). Genes regulated by *p53* show peaks of expression at every stage of the trajectory (Figure 3C). This is aligned with the known effect of 5-aza-CdR.

Additionally, we can also perform transcription factor enrichment analysis on these genes. (see online methods) This reveals several key regulators strongly associated with the inferred trajectories (Figure 3D, Supp Figure 6)

Amongst regulators of the cell cycle, the most significantly enriched one is *E2F4* with a p-value of 4x10-6, the gene enrichment also showed a significant p-value for the enrichment of *E2F* targets. Members of the *E2F* family are central regulators of cell cycle progression, particularly controlling the transition from G1 to S phase. Furthermore, they are known to interact with tumor suppressor proteins such as the Rb family. Consistent with prior studies, *E2F4* overexpression has been associated with colorectal tumor development, suppression of apoptosis, and increased proliferative capacity across multiple cancer types, thus acting as an oncogene ^32^. In addition, several members of the E2F family have already been identified to be connected to 5-aza-CdR in other cancers, thus suggesting a similar effect also in colorectal cancer^28,33,34^.

The transcription factors *JUND* (p-value 7x10-3) and *FOSL2* also emerge together. These two genes form the AP-1 complex, which has a regulatory role in colon cancer ^35^. *FOSL2* specifically has also been shown to promote metastasis in colon cancer ^36^.

Our study recovers known cancer-fighting mechanisms. *MYC* (p-value 6x10-4), *is* a well-established oncogene and major downstream effector of the WNT signaling pathway. In colorectal cancer specifically, loss of APC (which encodes a tumour suppressor protein) leads to activation of the WNT pathway^37^. The WNT pathway drives proliferation through *MYC*, which has been shown to be required for the majority of WNT target gene activation. In our data, *MYC* appears downregulated along the trajectories, consistent with its role as a therapeutic target and with previous reports showing that demethylating treatments can induce senescence through modulation of *MYC* activity. 5-aza-CdR has already been shown to induce senescence via reduction of *MYC* activity in Chronic Myelogenous Leukemia^38^. Our results suggest a similar effect in colorectal cancer. Our second most significantly enriched transcription factor is *CEBPD* (p-value 5x10-4), a known target of DNA demethylating agents. Indeed, *CEBPD* is frequently silenced by DNA methylation in cancer demethylating agents such as 5-aza-CdR have been shown to restore *CEBPD* expression and induce downstream apoptotic programs^39^. In hepatocellular carcinoma (liver cancer) 5-aza-CdR restored CEBPD expression, reducing tumor growth^40^. *FOXM1* (p-value 2x10-3) is generally involved in cell proliferation and is a well-established oncogenic factor promoting cell cycle progression. In colorectal cancer cells specifically, it has been shown to regulate metastasis through epithelial to mesenchymal transition and could constitute a potential therapeutic target^41^. In acute myeloid leukemia (AML) treatment with 5-Aza-2′-deoxycytidine increased the expression of miR-370, which has been shown to inhibit leukemic cell proliferation and induce senescence through targeting FOXM1^42^.

We identify several transcription factors that may represent previously unknown targets of demethylating treatment. *ID2 (*p-value 6x10-4) is associated with cell proliferation and cancer progression. *ID2* regulates cellular growth, senescence, differentiation, and apoptosis. The knockdown of *ID2* in HCT116 cells, as previously studied in mice, results in reduced proliferation and invasion of cancerous cells, leading to cell cycle arrest^43^. In parallel, *FLI1* (p-value 2x10-3) and *SPDEF* emerge as tumor suppressive regulators potentially reactivated by DNA demethylation. *FLI1* is known to be one of the most hypermethylated genes in colorectal cancer, with hypermethylation associated with poor prognosis, while its re-expression has been linked to reduced tumor growth and increased apoptosis. The reactivation of *FLI1* after demethylation may represent a previously unknown mechanism of the anti-cancer effects of 5-aza-CdR. Similarly, *SPDEF* has been shown to inhibit proliferation in colon cancer cell lines, including HCT116. Together, these findings suggest that, beyond known targets such as *CEBPD*, demethylating treatment may act through modulation of additional transcriptional programs involving *ID2, FLI1*, and *SPDEF*, which could contribute to limiting tumor growth and promoting anti-cancer cellular states following DNA demethylation.

### 4. FLOWSTATE reveals early establishment of signaling programs underlying motor neurons commitment and positioning

We investigate the differentiation of neuromesodermal progenitors into multiple fates in mice. Several key processes happen in the early stages, as it first captures the differentiation of neuromesodermal progenitors (NMPs) into neural and mesodermal lineages. This is followed by the differentiation of neural progenitors into three fates: floor plate (FP) cells, motor neurons (MNs), and V3 interneurons (V3). The determinants behind those different trajectories remain incompletely understood. To better understand this process we considered a dataset of mouse embryonic stem (ES) cells in the earlier stages of development^3^. The cells are sequenced every day between 3 and 8 days of development following a two-hour labeling period (Figure 4A).

**Figure 4.**
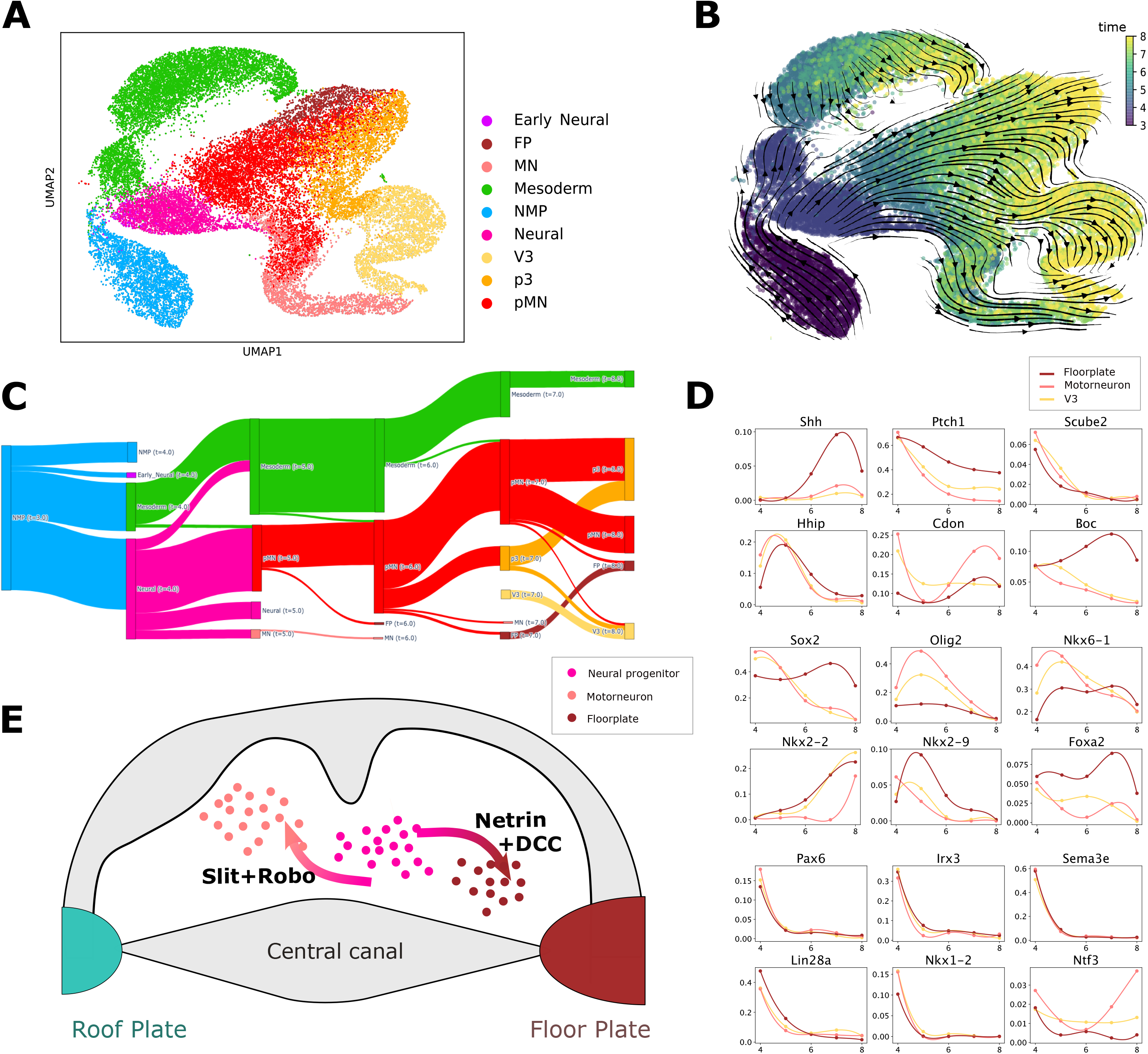
Trajectory inference on mouse embryonic stem cells differentiation. A. Umap with annotated cell types B.Velocity derived from the potential plotted on Umap with cells colored by time C. Cell type transition across time D. Mean expression of relevant genes across different fates E. Schematic representation of cell positioning in the ventral tube under the join effect of *Slit/Rob*o-mediated repulsion and *Netrin-1/DCC-*mediated attraction

After training FLOWSTATE, we assess the coherence of the inferred energy landscape. We assign a velocity to each cell, defined as the negative gradient of the potential. Visualizing this velocity field on a UMAP embedding reveals coherent flow across time points, with earlier cells being associated with higher potential (Figure 4B).

Next we computed cell type transitions across time points as described in the benchmarking section. Between day 3 and day 4, 49% of neuromesodermal progenitors (NMPs) transition to an early neural state, while 33% transition to mesoderm (Figure 4C). The mesoderm population remains largely stable throughout the remainder of the time course, with 82% not undergoing further transitions to other cell types and at least 92% remaining stable between consecutive time points. The remaining 12% are mostly classified as transitioning into pMN, which is likely an artifact due to the very high proportion of pMN cells in the dataset. Neural progenitors subsequently evolve into three final states: motor neurons (MN) via pMN, and VP and F3 via p3. We observe that the MN fate emerges earlier than the VP/F3 states. Indeed, a stable pMN population appears as early as day 5, whereas p3, V3, and FP populations only emerge later. The initial branching into MN versus FP/V3, followed by a secondary split between FP and V3, is consistent with observations from the original paper as well as previous studies^44,45^.

Finally, we focus in particular on the fate of early neural cells at day 4, which can differentiate into three lineages: Motor Neurons (MNs), V3, and Floor Plates (FPs). To analyze the evolution of gene expression along these trajectories we plot the average expression value of each gene in each trajectory across time (see online methods). As shown in Figure 4D we recover in this way trends consistent with those reported in the original study. Some differences are to be noted, however. For the few genes where our predicted trends are different from the original paper, our findings are more biologically coherent. For instance, *PAX6* is known to be highly expressed in cells that commit to the MN fate. In our analysis, we correctly infer PAX6 to be higher expressed in cells that commit to the MN fate with respect to V3 and FP for early phases of the trajectory^46^. On the contrary, the original study reports a higher expression of PAX6 in the V3 fate, suggesting that our inferred trajectory is more biologically coherent. Similarly, we capture an increase in the expression of *NTF3* along the MN differentiation trajectory, while the original study predicts a decrease of this gene towards the MN fate and no difference among the three commitment states. Also in this case, our prediction fits best the known biology of the system^47^.

We next perform unsupervised discovery of gene expression trends as we have done previously. To understand which genes are determinants of differentiation, we perform differential gene analysis across the three trajectories (see online methods). Among the genes more highly expressed in cells that commit to the MN fate, we notably find *Robo2* and *Slit3*, while *Ntn1* is significantly more expressed in cells that commit to the FP fate. Previous findings show that motor neurons positioning is regulated by the balance between *Slit/Rob*o-mediated repulsion and *Netrin-1/DCC-*mediated attraction^48^. In this context, *Slit* ligands produced by the floor plate repel motor neurons via *Robo* receptors, preventing their invasion of the ventral midline, whereas floor plate-derived *Netrin-1* attracts motor neurons via the *DCC* receptor, drawing them toward the floor plate (Figure 4E). The enrichment of *Robo2* expression in MN-committed cells is consistent with the established role of Robo receptors in mediating Slit-dependent repulsion of motor neurons from the floor plate.

Furthermore, the expression of *Slit3* in MN-committed cells is consistent with previous reports that motor neurons themselves can provide Slit-mediated signals in an autocrine and/or paracrine manner. Such signaling has been proposed to activate *Robo* receptors and limit the attractive effects of *Netrin/DCC* signaling, thereby preventing inappropriate migration toward the ventral midline. Our observed enrichment of both *Slit3* and *Robo2* in MN-committed cells between days 3 and 8, suggest that components of this signaling axis might be established early during motor neurons commitment, whereas previous studies have primarily described Slit–Robo signaling in later-stage differentiating motor neurons during migration and axon guidance^49^.

## Discussion

Recent advances in single-cell technologies have enabled a deeper understanding of cellular dynamics through metabolic RNA labeling. These approaches allow the distinction between nascent and pre-existing RNA, providing direct access to transcriptional dynamics at the single-cell level. Fully leveraging this information requires trajectory inference methods specifically designed to incorporate the additional information provided by labeled RNA data.

Here, we introduced FLOWSTATE, a trajectory inference method that directly integrates dynamical information derived from labeled RNA into its learning objective. We benchmarked FLOWSTATE across three datasets, evaluating its ability to accurately predict unseen snapshots. Our results demonstrate consistent improvements over methods that do not incorporate labeled RNA information. Furthermore, FLOWSTATE achieves more accurate recovery of cell-type transitions, highlighting its ability to capture more biologically meaningful dynamics.

Although demonstrated using metabolic RNA labeling data, the method is agnostic to the source of dynamical information and can incorporate any experimental or computational estimate of cellular dynamics. FLOWSTATE therefore provides a general framework for integrating emerging dynamic single-cell measurements from any future technology into trajectory inference models, whether derived from alternative RNA-labeling methods or from entirely different technologies.

We then applied FLOWSTATE to two datasets and performed unsupervised discovery of driver genes and transcription factors. The first dataset consists of colorectal cancer cells treated with a DNA demethylating agent, while the second captures neural progenitors differentiating into multiple neural lineages. In both systems, cells undergo rapid transcriptional changes, which are better captured by labeled RNA than by traditional single-cell sequencing approaches. In colorectal cancer HCT116 cells, FLOWSTATE recovers well studied effects of the 5-aza-CdR treatment and identifies *FLI1* and as potential new treatment targets. In mouse embryonic stem cell differentiation, we find that components of the *Slit/Robo* signaling axis are enriched in MN-committed cells, suggesting that this known pathway might be established during early motor neurons commitment, prior to the later stages of migration and axon guidance in which it has traditionally been studied.

However, several limitations remain. Experimental protocols and labeling strategies vary substantially across datasets, requiring different preprocessing. In benchmarking, the lack of ground-truth pairings between cells across time points restricts evaluation to distribution-level comparisons at the final time point. While metrics based on cell-type transitions provide additional insight into the correctness of inferred pairings, they do not fully resolve this limitation.

Our modeling framework is based on representing cellular evolution as the minimization of a potential energy, rooted in the concept of the Waddington landscape. This approach provides interpretability, as the influence of individual genes across different regions of the potential can be analyzed. However, it also imposes constraints on the types of dynamics that can be modeled. In particular, the potential depends only on gene expression, thereby neglecting cell–cell interactions. Moreover, because cells evolve toward lower potential states, cyclic or non-equilibrium processes cannot be accurately captured within this framework.

Future work could address these limitations by extending the model to incorporate more flexible energy functions or interaction mechanisms. For example, incorporating quadratic energy terms or attention-based interactions between cells could enable the modeling of more complex dynamics, including cyclic behaviors and collective effects. Such developments represent promising directions for improving trajectory inference in dynamic single-cell systems.

## Method section

### Data collection

#### Lung cancer cells

*benchmark*. The cells are sequenced after 0h, 2h, 6h, 8h, and 10h of treatment with dexamethasone (DEX) . The cells are labeled with 4sU for 2h before the sequencing. The sci-fate technique is used for labeling and sequencing. We use the first 4 time points as a train set and the last one as a test set. We have 6673 cells after preprocessing.

#### Excitatory neuron cells

*benchmark*. The cells are neurons being activated for different durations: 15 min, 30 min, 60min and 120min. The data used is shown in Figure 3 of the original publication. The cells are labeled with 4sU for 2h before the sequencing. The scNT-seq technique is used for labelling and sequencing. We use the first 3 time points as a training set and the last one as a test set. We have 3060 cells after preprocessing.

#### Embryonic stem cells

*benchmark and case study*. The cells are differentiating into neural tube identities. The cells were sequenced daily from day 3 to day 8. The cells are labeled with 4sU for 2h before the sequencing. The scNT-seq technique is used for labeling and sequencing. For the benchmarking our test set is made of the last time point of replicate 1, as well as the full replicate 2 and replicate 3. For the case study, we use all three replicates. We have 33527 cells after preprocessing.

#### HCT116 colorectal cancer cells

*case study*. The cells are being treated with 5-AZA-CdR and sequenced after 0, 1, 2, and 3 days of treatment. The cells are labeled with 4sU for two hours before sequencing. The well-TEMP-seq labeling and sequencing technique is used. The data used is shown in Figure 5 of the original publication. We have 3213 cells after preprocessing.

### Notations

We have access to the cell distributions at time points *t*_1_ < … <*t*_*k*_ <… <*t*_*K*_. For any time point *t*, we denote by 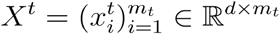 the corresponding dataset, where *m*_*t*_ is the number of cells and *d* is the dimension of the representation space. The vector 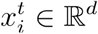 represents the gene expression profile of the *i*-th cell belonging to time *t*, either in the gene space or in a reduced PCA space. We also observe labeled (new) RNA, denoted by *n*. This corresponds to RNA transcripts produced during a labeling interval of duration *τ* before sequencing. Similarly, we define 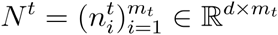 For every cell *i* and gene *g*, we have *n*_*i,g*_ < *x*_*i,g*_ The vector *n*_*i*_ ∈ ℝ^*d*^ represents the newly synthesized RNA of the *i*-th cell. We define the empirical distribution associated with time point *t* as 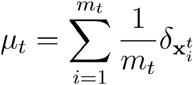 For a function *f* : ℝ^*d*^ → ℝ^*d*^, we define the pushforward measure as 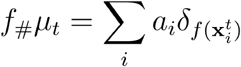.

### Inference of Cell Dynamics from Labeled RNA

For any gene, the evolution of the quantity of total and new RNA is described by the following equations:

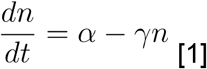

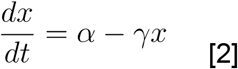

where *n* and *x* denote the quantities of labeled and total RNA, respectively, and where *α* and *γ* are the gene-specific transcription and degradation rates.

#### Matching a target distribution

In the first framework, for any time point *t*_*k*_ (except the first one), the labeling time *τ*_*k*_ satisfies *τ*_*k*_ = *t*_*k*_ − *t*_*k*−*l*_ for some 0 < *l* < *k*.

We solve Equations [1] and [2] with initial conditions n(*t*_*k*−*l*_) = 0 and 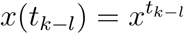 This gives

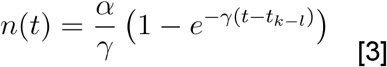

and

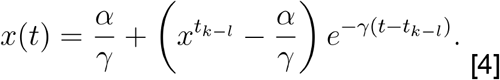

Therefore,

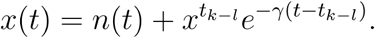

For any cell *i* sampled at time point *t*_*k*_, the estimated expression at time *t*_*k*−*l*_ is

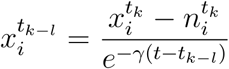

For a given degradation rate, we define

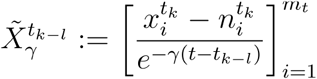

This corresponds to an estimate of the distribution at the time point *t*_*k*−*l*_.

We estimate the degradation rate by minimizing the squared 2-Wasserstein distance between the estimated and observed distributions plus a regularizing term of the total mass of the cell we use α = 0.1:

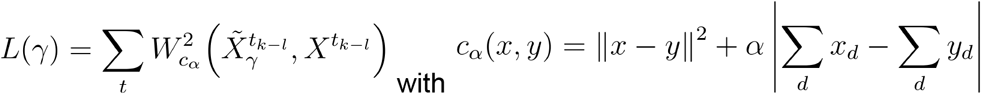

We directly estimate 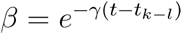 for each gene. We also replace *n* by *n/δ* with *δ* being the detection rate of the labeled RNA (what fraction of transcripts are being labeled). We use *δ* = 0.83.

For every cell belonging to any time point we have access to the gene expression of the cell as well as an estimation of the position of the cell at time 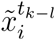. We can define 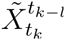 the approximated state of the cells from *t*_*k*_ at time *t*_*k*−*l*_. We check that the corrected distributions are well aligned and that most of the estimated degradation rates in between each time interval are in the coherent biological range [0,1] (see Supp Figure 1). Our method is sufficiently general to be applied to any dataset where we have a matching between the beginning of the labelling time and one of the previous time points regardless. It can adapt to the case where we have a one to one pairing between time points or one time point can correspond to the beginning of the labeling for several of the later times.

#### Computing a local velocity

This framework follows the strategy proposed in previous works^11,50^ . Here, the labeling time *τ* is assumed to be short compared to the interval between two sampled time points.

From Equation [3], we derive the transcription rate

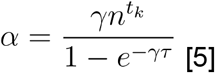

The resulting velocity is

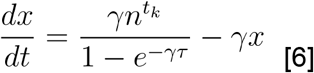

We estimate *γ* assuming steady-state dynamics. n = *kx*_*ss*_ and 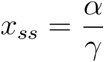 Substituting into Equation [5] gives

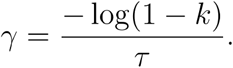

We estimate *k* using extreme-value regression between new and total RNA counts and compute the corresponding degradation rate *γ*. Finally, Equation [6] is used to estimate the instantaneous velocity *v*_*i*_ of each cell. The velocity vectors are rescaled such that, at each time point, their mean magnitude matches the average transport distance to the next time point, where pairwise euclidean distances are weighted by the optimal transport coupling.

### Preprocessing

All datasets were handled as anndata object ^51^ for every dataset, we performed the following preprocessing steps on the total RNA.

#### Cell and gene quality control

Using Scanpy’s ^52^ *sc*.*pp*.*filter_cells, sc*.*pp*.*filter_genes*. We removed low-quality cells and genes expressed in too few cells. We used the threshold used in the original publication of each dataset.

#### Normalization and log transform

We applied Scanpy’s *sc*.*pp*.*normalize_total* and *sc*.*pp*.*log1*

#### Selecting the highly variable genes

We applied Scanpy’s *scanpy*.*pp*.*highly_variable_genes*. When multiple replicates (embryonic stem cell dataset) are available, we select 3000 genes and keep the genes that are highly variable in all replicates. In the set up where we match a target distribution the matching is done based on the 4000 most variable genes. Only genes whose estimated degradation rate in between every time interval are in the coherent biological range [0,1] are kept.

#### Scaling

Using Scanpy’s sc.pp.scale, we scale data to unit variance and zero mean.

#### Dimensionality reduction

Using Scanpy’s *sc*.*tl*.*pca*, we applied Principal Component Analysis (PCA) to reduce the data to 15 dimensions for data set 1 and 2 and 20 dimensions for data set 3 and 4.

#### Visualization

Using Scanpy’s *sc*.*tl*.*umap*, we applied UMAP to project the batch-corrected data into two dimensions.

#### Preprocessing the labeled RNA

Preprocessing when matching a target distribution (lung cancer dataset and excitatory neuron dataset): Cell and gene quality control is performed based on the total RNA only. After the estimated distributions at past state have been computed, normalization log transform and scaling is applied and PCA projection is applied globally. Preprocessing when estimating a local velocity (embryonic stem cells and colorectal cancer cells : for the normalization, log transform, and scaling, we project the velocity in the new space using the Jacobian of the transformation. applied to the total RNA. We then project the velocity into the PCA space using the PCs computed on the total RNA.

### Wasserstein gradient flow learning

#### Optimal transport

Let 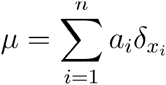 and 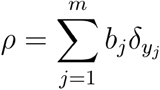 be two discrete probability distributions on *R*^*d*^.

The squared 2-Wasserstein distance^53^ between μ and ρ is defined as

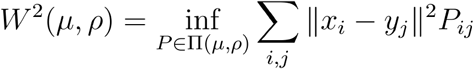

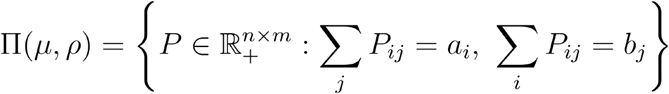 denotes the set of transport couplings.

This problem can be regularized with an entropy term. The entropy-regularized optimal transport ^54^ “distance” is

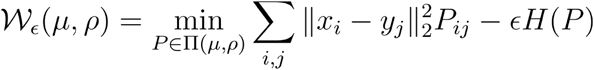

where 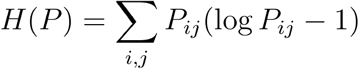 . Since 𝒲_*ϵ*_(*μ, μ*) ≠ 0 we use the debiased Sinkhorn divergence^25^ instead

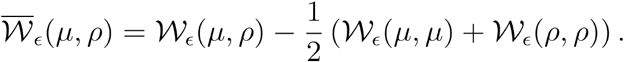

#### Wasserstein gradient flow

We model the evolution of *μ*_*t*_ as a Wasserstein gradient flow associated with a potential energy.

For a single cell *x*, the Euclidean gradient flow satisfies 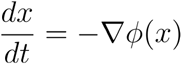, which is the continuous counterpart of gradient descent.

Wasserstein gradient flows^55^ extend this formulation to probability measures. We say that μ_*t*_ follows a Wasserstein gradient flow associated with the free energy functional ℱ : μ ↦ *∫ ϕ*(*x*) *dμ* (*x*) ∈ ℝ if its density satisfies the continuity equation :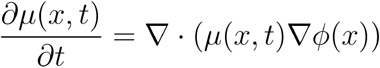.

In the case of discrete distributions this is equivalent to each particle following the above mentioned euclidean gradient flow.

We want to learn a neural network *ϕ*_*θ*_ from the observed snapshots 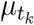 such that for given parameters *θ* and an initial population 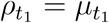, the predicted populations 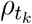 are close to the observed snapshots 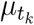.

In order to do our prediction, we use the forward Euler discretization scheme 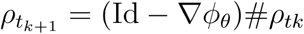 . Meaning that for each cell 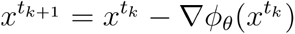 for each cell *x*. We only do one step between each time point. We use teacher forcing, meaning we predict 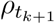 from 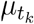 rather than 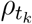. We define the same equation in parallel over a set of cells *X*^*t*^ as

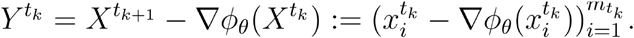

The predicted distribution associated with time point *t*_*k*−1_ is 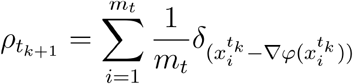.

In practice proposed to parameterize the potential *ϕ* by a simple one-hidden-layer MLP *ϕ*_*θ*_ and we minimize the loss *L*(*θ*) over the parameters of the neural network θ

### Loss function

#### Loss function when matching a target distribution

In this setup, we can estimate the state of each observed cell at an earlier time point. For instance, in the lung cancer dataset (dataset 1), for every cell at time *t*_*k*_ we have an approximation of its state at the previous time point *t*_*k*−1_. In the neuron (dataset 2), for every cell we have an approximation of its state at time *t*_0_ (Fig.2A).

Let us define the recursive prediction operator *Φ*_θ_,(*X, n*_steps_) which takes as input a cell distribution and a number of steps *n*_steps_, and outputs the predicted cell states after recursively applying the learned dynamics *n*_steps_ times.

To compute *Φ*_θ_,(*X, n*_steps_) we define *Y*^0^ = *X* and for *i* ∈ [0, *n*_steps_−1], *Y*^*i*+1^ = *Y*^*i*^ − ∇*ϕ*_*θ*_ (*Y*^*i*^).

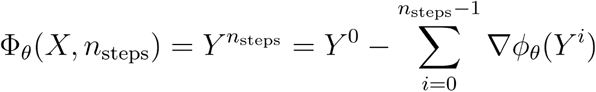

Rigorously, for any *k*, there exists a *σ* (*k*) < *k* such that we can approximate the state of the cells from *t*_*k*_ at the earlier time 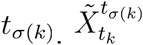. denotes the estimated state at time *t*_*σ* (*k*)_ of the cells belonging to time *t*_*k*_ (see section Inference of Cell Dynamics from Labeled RNA/Matching a target distribution)

In the lung cancer dataset *∀k, σ* (*k*) = k − 1 while in the neuron dataset *∀k, σ* (*k*) = 0.

We recursively apply the learned vector field − ∇*ϕ*_*θ*_to this estimate to evolve the cell state forward in time until we reach time 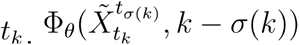. is a prediction of the distribution at time *t*_*k*_, where each predicted cell is paired with a cell from 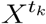. For simplicity we will write 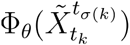 instead of 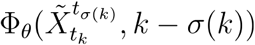

We minimize the following loss:

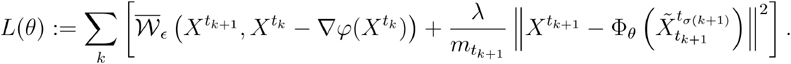

Indeed, cells from 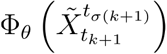 are directly paired with cells from 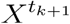, and therefore no optimal transport step is required to compare the two distributions. We divide by 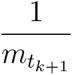 to make the scale of this term comparable with the sinkhorn term. The term 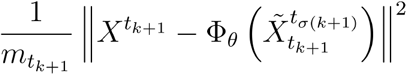 can be interpreted as a squared 2-Wasserstein cost computed under the fixed coupling *P* = Id induced by the identity pairing between cells.

In the second term of the loss, we can rewrite :

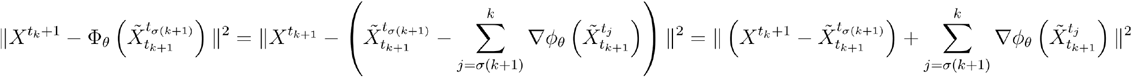

Here for 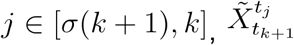 denotes 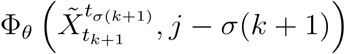, the intermediary prediction made while computing *Φ*_*θ*_. Those correspond to the *Y*^*k*^ mentioned above in the recursive computation of Φ _*θ*_. Since we know that the cells of 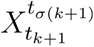, evolve into the cells at 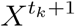 we can view 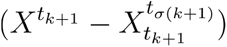 as an estimate of the displacement of those cells between *σ*(*k*+1) and *t*_*k*+1_

#### Loss function when computing a local velocity

For every cell belonging to any time point we have 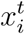 the gene expression of the cell and 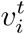 estimate of the instantaneous velocity of the cell. This is the case in the embryonic stem cell dataset (dataset 3) as well as in the colorectal cancer cells dataset. Let us define 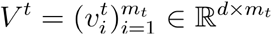 the estimate of the instantaneous velocity of cells at time *t*

We minimize the following loss:

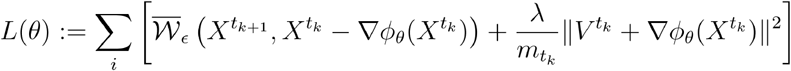

#### General formulation of the loss

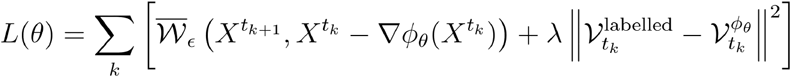

When matching a target distribution, we have :

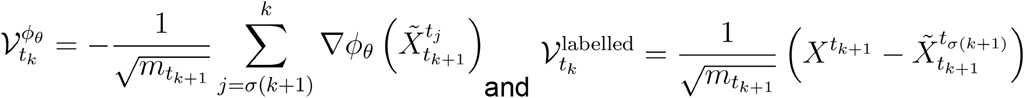

In this setting, the term aligns the *ℓ*^2^ accumulated trajectory predicted by the learned Wasserstein gradient flow with trajectory information inferred from labeled RNA measurements.

When computing a local velocity, we have :

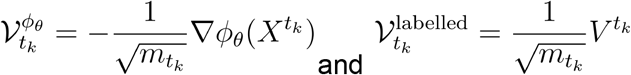

In this setting, the term aligns the instantaneous velocity predicted by the learned Wasserstein gradient flow with instantaneous velocity estimates derived from labeled RNA measurements.

#### Choice of λ parameter

Here *λ* > 0 is a parameter to be tuned that characterized the weight we give to the ℓ^2^ loss and the coupling information with respect to the OT-loss. When we simply train using the sinkhorn divergence. We tested the value of λ ∈ 0,0.1, 0.25, 0.5, 0.75, 1.0, 1.25. We reported results for *λ =* 0.25 in the benchmarking as this was the best of second best performing values in all cases.

### Computational Optimal Transport

**OTT solvers**. We use the OTT package ^56^ to solve OT problems. In particular, we rely on the Sinkhorn solver. We set the regularization weight to *ϵ* = 0.01

### Neural network training

#### Neural network architecture

We implemented a Multi-Layer Perceptron (MLP) using Flax ^57^. The network is composed of two hidden layers of dimension 128 with GeLU activation functions. We do not include a bias term in the final layer as it would not affect the values of ∇*ϕ*_*θ*_. Additionally, we used a smooth GeLU activation function over the standard ReLU.

#### Data loading

For each time point described, we split the cells into 75% training samples and 25% validation samples. For both training and validation, batches were constructed by uniformly sampling 15% of cells in each time point without replacement. Validation was performed once every four iterations.

#### Optimizer

We used Optax’s implementation of the AdamW optimizer. The learning rate was scheduled using Optax’s cosine decay scheduler, with an initial value of 1e-3 over 10,000 decay steps.

#### Early Stopping

The maximum number of iterations is set to 15000, but training was stopped early if the validation loss did not improve for 150 consecutive iterations. We kept the model weights corresponding to the lowest validation loss.

#### Seeds

We ran every experiment with 5 seeds 1, 2, 3, 4, 5. The seed reproducibly determines the train/validation split and weight initialization. For the two case studies “FLOWSTATE highlights the downstream effects of 5-aza-CdR and identifies potential treatment targets in colorectal cancer HCT116 cells” and “FLOWSTATE reveals early establishment of signaling programs underlying motor neuron commitment and positioning” the experiment corresponds to seed 1. For the visualization of cell type transition in Fig 2. the seed displayed is 4.

### Benchmarking scores

#### Wasserstein distance

We compute the 2-Wasserstein distance (the unregularized optimal transport) between the predicted and ground truth distribution for the last time point kept out during the training. The Wasserstein distance, according to the squared Euclidean distance, is computed using the OTT package ^56^

#### Sinkhorn divergence

We compute the Sinkhorn divergence between the predicted and ground truth distribution for the last time point kept out during the training. We use *ϵ* = 0.01 and also use the OTT package.

#### CBD Score

We compute cross-boundary direction (CBD) correctness using the implementation from ^58^. CBD quantifies the accuracy of predicted transitions from a source cluster to a target cluster using boundary cells, defined as cells whose neighbors belong to the opposite cluster. For each cell, CBD is computed as the average cosine similarity between its inferred velocity vector (−∇*ϕ*(*x*)) and the displacement vectors toward neighboring cells in the target cluster.

#### Cell type transition accuracy

After predicting each cell’s future states, we assess the accuracy of our cell type transition matrix by comparing predicted transitions between cell types to a set of biologically allowed transitions. In practice, we calculate the fraction of total transition probability mass that corresponds to allowed (correct) transitions.

### Running prescient

We benchmarked FLOWSTATE against PRESCIENT, which also leverages optimal transport (OT) to learn a gene expression potential within a causal model of differentiation. We used the authors’ open-source implementation (https://github.com/gifford-lab/prescient/). We ran PRESCIENT with the default settings, except for the train_sd parameter. The default value of 0.5, which determines the magnitude of stochastic noise in the differentiation dynamics, was too large for the scale of gene expression in the benchmark datasets. To ensure a fair comparison, we did a grid search and we adjusted this parameter to train_sd = 0.005 (see Supp Figure 5).

### Computational growth rate

#### Weight of marginals

When comparing a prediction 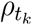 to the reference snapshot 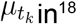 in^18^ Yeo et al. proposes setting the weights *b*_*j*_ of the prediction 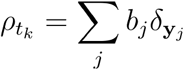 proportionally to a computationally derived growth rate. This allows us to take into account the fact that some cells should be matched to a larger number of descendants by assigning a higher growth rate to them. This approach was introduced by Schiebinger^14^ and reimplemented in MOSCOT^16^. Here, we follow MOSCOT’s implementation.

#### Proliferation and apoptosis

The computation of growth rates is based on cell-wise proliferation and apoptosis scores. The proliferation and apoptosis scores were computed with Scanpy’s sc.tl.score_genes function using gene sets curated from the literature. The scores are calculated on raw counts after quality filtering.

#### Calculating the growth rate

For a given cell *x*_*i*_, let prol_*i*_ ∈ ℝ and apo_*i*_ ∈ ℝ be the proliferation score and the apoptosis score ^16^, respectively. The birth rate and *β*_*i*_∈ ℝ death rate *γ*_*i*_∈ ℝ are defined as

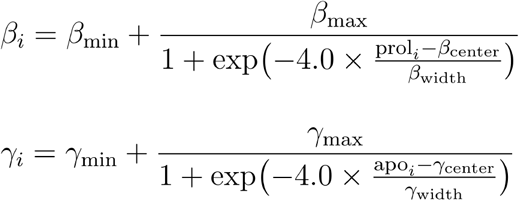

We used MOSCOT’s default parameters *β*_min_ *= γ*_min_=0.3, *β*_max_ *= γ*_max_= 1.7, *β*_center_ *=* 0.25, *β*_width_= 0.5, *γ*_center_ =0.1, *γ*_width_ *=* 0.2.

The growth rate of the cell is then defined as *g*_*i*_ = expt (Δ*t* × (*β*_*i*_ − *γ*_*i*_)) where Δ*t* denotes the time difference between populations.

The growth rates are normalized to obtain weights *a*_*i*_ that sum to 1: 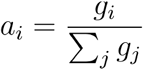,

In this formulation, Δ_*t*_ acts as the inverse temperature of the softmax. Larger values of Δ_*t*_ produce a sharper distribution. Since time points in our experiments were evenly spaced, we use Δ_*t*_ *=* 1. We used growth rate in the second case study “FLOWSTATE reveals early establishment of signaling programs underlying motor neurons commitment and positioning” (see Supp Figure 7). In the benchmark the marginals were kept uniform to focus on the quality of the models.

### Biological analysis

#### Cell type transition matrix and prediction of each cell’s future cell type

First, each cell is pushed forward in time by the model, which gives a predicted future position. The cell *x* moves to the position x −∇*ϕ*_*θ*_(*x*). We use a k-nearest neighbors algorithm trained on the entire dataset to determine the cell type of the predicted future state of the cell. The k-NN is implemented using scikit-learn ^59^ and uses the 20 nearest neighbors in the PCA space.

#### Transcription factor enrichment

We perform transcription factor–target enrichment using the CollecTRI resource, which compiles curated regulatory interactions between transcription factors and their target genes in humans. First, we restrict the interaction network to targets present in the dataset and construct, for each transcription factor, a binary indicator over genes specifying whether each gene is a known target. We then compare the distribution of regression-derived scores (reflecting association with the inferred potential) between the set of *n*_1_ target genes and the complementary set of *n*_2_ non-target genes. For each transcription factor, we apply a one-sided Wilcoxon rank-sum test to assess whether its targets exhibit significantly higher regression scores than non-targets. Because the number of annotated targets varies across transcription factors, *n*_1_ differs while *n*_1_ + *n*_2_ equals the total number of genes considered. Transcription factors are subsequently ranked by their P values, and the top candidates are visualized the TF ranked by log_10_(*p*_*value*_), highlighting regulators whose targets are most strongly associated with the inferred potential.

#### Gene set and pathway enrichment analysis

We perform functional enrichment analysis on the list of genes ranked by their correlation with the inferred potential using gseapy’s Enrichr function. The input gene list corresponds to genes most strongly associated with the potential. The databases of gene sets and pathways used are Gene Ontology Biological Process, Gene Ontology Molecular Function, and Gene Ontology Cellular Component, MSigDB Hallmark, KEGG (2021 Human), Reactome (2024), and WikiPathways (2024 Human) databases.

#### Fate classification

To map cells across time points and get a tractable evolution starting from the initial population, we compute the optimal transport plan between predicted positions and observed cells at the next time point. This transport plan defines a probability distribution mapping each source cell to cells at the subsequent time point, describing how the mass of each cell is distributed over its potential descendants. We propagate this probability mass forward across successive time points, yielding, for each starting cell, a distribution over cells reachable at later stages. These cell-level probabilities are then aggregated by terminal identity, assigning each starting cell a probability of ending in each major fate. The most likely final identity is used to classify cells into three branches (FP, MN, and V3). In this way, each trajectory is defined not only by a terminal label but also by a continuous set of cells connected through propagated probability flow from the initial neural population.

## Data availability

### Cell lines

The lung cancer dataset can be downloaded in raw and processed forms from the NCBI Gene Expression Omnibus (GEO) the GEO accession number is GSE131351 the data can be found at https://www.ncbi.nlm.nih.gov/geo/query/acc.cgi?acc=GSE131351

The neuron dataset can be downloaded at https://www.dropbox.com/s/5wk3q2xhgqai2xq/neuron_labeling.h5ad?dl=1

The embryonic stem cell dataset is accessible on GEO with the number GSE236520 and the data is available at https://www.ncbi.nlm.nih.gov/geo/query/acc.cgi?acc=GSE236520. The colorectal cancer cells dataset raw data files are available at NCBI Gene Expression Omnibus (GEO) with the number GSE194357 (https://www.ncbi.nlm.nih.gov/geo/query/acc.cgi?acc=GSE194357).

## Code availability

*Package*. The Python package for FLOWSTATE is hosted at https://github.com/cantinilab/FLOWSTATE. Code to reproduce the experiments and figures is available at https://github.com/cantinilab/FLOWSTATE_reproducibility

## Acknowledgements

The project leading to this manuscript has received funding from the European Union, European Research Council StG and MULTIview-CELL (101115618, to L.C.). In addition, this work was supported by a government grant managed by the Agence Nationale de la Recherche under the France 2030 program, with the reference numbers ANR-24-EXCI-0001,ANR-24-EXCI-0002, ANR-24-EXCI-0003, ANR-24-EXCI-0004, ANR-24-EXCI-0005 (to L.C.). This work was performed using HPC resources from GENCI–IDRIS (Grant 2021-AD011013214). We acknowledge the help of the HPC Core Facility of the Institut Pasteur and Deborah Philipps for the administrative support. The work of G. Peyre was supported by the French government under the management of Agence Nationale de la Recherche as part of the “Investissements d’avenir” program, reference ANR-19-P3IA-0001 (PRAIRIE 3IA Institute) and by the European Research Council (ERC project WOLF).

## References

1. Tritschler, S. et al. Concepts and limitations for learning developmental trajectories from single cell genomics. Development 146, dev170506 (2019).

2. Erhard, F. et al. Time-resolved single-cell RNA-seq using metabolic RNA labelling. Nat. Rev. Methods Primer 2, 77 (2022).

3. Maizels, R. J., Snell, D. M. & Briscoe, J. Reconstructing developmental trajectories using latent dynamical systems and time-resolved transcriptomics. Cell Syst. 15, 411-424.e9 (2024).

4. Lin, S. et al. Well-TEMP-seq as a microwell-based strategy for massively parallel profiling of single-cell temporal RNA dynamics. Nat. Commun. 14, 1272 (2023).

5. Saelens, W., Cannoodt, R., Todorov, H. & Saeys, Y. A comparison of single-cell trajectory inference methods. Nat. Biotechnol. 37, 547–554 (2019).

6. Weiler, P., Van den Berge, K., Street, K. & Tiberi, S. A Guide to Trajectory Inference and RNA Velocity. in Single Cell Transcriptomics: Methods and Protocols (eds Calogero, R. & Benes, V.) 269–292 (Springer US, New York, NY, 2023). doi:10.1007/978-1-0716-2756-3_14.

7. Street, K. et al. Slingshot: cell lineage and pseudotime inference for single-cell transcriptomics. BMC Genomics 19, 477 (2018).

8. Ji, Z. & Ji, H. TSCAN: Pseudo-time reconstruction and evaluation in single-cell RNA-seq analysis. Nucleic Acids Res. 44, e117 (2016).

9. Trapnell, C. et al. The dynamics and regulators of cell fate decisions are revealed by pseudotemporal ordering of single cells. Nat. Biotechnol. 32, 381–386 (2014).

10. La Manno, G. et al. RNA velocity of single cells. Nature 560, 494–498 (2018).

11. Bergen, V., Lange, M., Peidli, S., Wolf, F. A. & Theis, F. J. Generalizing RNA velocity to transient cell states through dynamical modeling. Nat. Biotechnol. 38, 1408–1414 (2020).

12. Li, C., Virgilio, M. C., Collins, K. L. & Welch, J. D. Multi-omic single-cell velocity models epigenome-transcriptome interactions and improves cell fate prediction. Nat. Biotechnol. 41, 387–398 (2023).

13. Wang, W. et al. RegVelo: Gene-regulatory-informed dynamics of single cells. Cell 0, (2026).

14. Schiebinger, G. et al. Optimal-Transport Analysis of Single-Cell Gene Expression Identifies Developmental Trajectories in Reprogramming. Cell 176, 928-943.e22 (2019).

15. Tong, A., Huang, J., Wolf, G., Dijk, D. van & Krishnaswamy, S. TrajectoryNet: A Dynamic Optimal Transport Network for Modeling Cellular Dynamics. Preprint at 10.48550/arXiv.2002.04461 (2020).

16. Klein, D. et al. Mapping cells through time and space with moscot. Nature 638, 1065–1075 (2025).

17. Bunne, C., Papaxanthos, L., Krause, A. & Cuturi, M. Proximal Optimal Transport Modeling of Population Dynamics. in Proceedings of The 25th International Conference on Artificial Intelligence and Statistics 6511–6528 (PMLR, 2022).

18. Yeo, G. H. T., Saksena, S. D. & Gifford, D. K. Generative modeling of single-cell time series with PRESCIENT enables prediction of cell trajectories with interventions. Nat. Commun. 12, 3222 (2021).

19. Hashimoto, T., Gifford, D. & Jaakkola, T. Learning Population-Level Diffusions with Generative RNNs. in Proceedings of The 33rd International Conference on Machine Learning 2417–2426 (PMLR, 2016).

20. Huizing, G.-J. et al. STORIES: learning cell fate landscapes from spatial transcriptomics using optimal transport. Nat. Methods 1–10 (2025).

21. Peng, Q., Zhou, P. & Li, T. stVCR: spatiotemporal dynamics of single cells. Nat. Methods 23, 542–553 (2026).

22. Allen, M. Compelled by the Diagram: Thinking through C. H. Waddington’s Epigenetic Landscape. Contemp. Hist. Presence Vis. Cult. 4, 119–142 (2015).

23. Cao, J., Zhou, W., Steemers, F., Trapnell, C. & Shendure, J. Sci-fate characterizes the dynamics of gene expression in single cells. Nat. Biotechnol. 38, 980–988 (2020).

24. Qiu, Q. et al. Massively parallel and time-resolved RNA sequencing in single cells with scNT-seq. Nat. Methods 17, 991–1001 (2020).

25. Genevay, A., Peyre, G. & Cuturi, M. Learning Generative Models with Sinkhorn Divergences. in Proceedings of the Twenty-First International Conference on Artificial Intelligence and Statistics 1608–1617 (PMLR, 2018).

26. I, V. et al. The scverse project provides a computational ecosystem for single-cell omics data analysis. Nat. Biotechnol. 41, (2023).

27. Jueliger, S., Taverna, P., Re, O. L. & Vinciguerra, M. Senescence Induced by DNA Demethylating Drugs to Treat Solid Tumors. in Handbook of Immunosenescence 1–30 (Springer, Cham, 2018). doi:10.1007/978-3-319-64597-1_166-1.

28. Momparler, R. L. Epigenetic therapy of cancer with 5-aza-2’-deoxycytidine (decitabine). Semin. Oncol. 32, 443–451 (2005).

29. Jones, P. A. & Baylin, S. B. The fundamental role of epigenetic events in cancer. Nat. Rev. Genet. 3, 415–428 (2002).

30. Van Tongelen, A., Loriot, A. & De Smet, C. Oncogenic roles of DNA hypomethylation through the activation of cancer-germline genes. Cancer Lett. 396, 130–137 (2017).

31. Virtanen, P. et al. SciPy 1.0: fundamental algorithms for scientific computing in Python. Nat. Methods 17, 261–272 (2020).

32. Mady, H. H., Hasso, S. & Melhem, M. F. Expression of E2F-4 gene in colorectal adenocarcinoma and corresponding covering mucosa: an immunohistochemistry, image analysis, and immunoblot study. Appl. Immunohistochem. Mol. Morphol. AIMM 10, 225–230 (2002).

33. Tonon, F. et al. 5-Azacytidine Downregulates the Proliferation and Migration of Hepatocellular Carcinoma Cells In Vitro and In Vivo by Targeting miR-139-5p/ROCK2 Pathway. Cancers 14, 1630 (2022).

34. Wang, W. et al. E2F2(E2F transcription factor 2) as a potential therapeutic target in meibomian gland carcinoma: evidence from functional and epigenetic studies. BMC Cancer 25, 880 (2025).

35. Ashida, R. et al. AP-1 and colorectal cancer. InflammoPharmacology 13, 113–125 (2005).

36. Li, S., Fang, X.-D.Wang, X.-Y. & Fei, B.-Y. Fos-like antigen 2 (FOSL2) promotes metastasis in colon cancer. Exp. Cell Res. 373, 57–61 (2018).

37. Sansom, O. J. et al. Myc deletion rescues Apc deficiency in the small intestine. Nature 446, 676–679 (2007).

38. Grandjenette, C. et al. 5-aza-2’-deoxycytidine-mediated c-myc Down-regulation triggers telomere-dependent senescence by regulating human telomerase reverse transcriptase in chronic myeloid leukemia. Neoplasia 16, 511–528 (2014).

39. Hartl, L., Duitman, J., Bijlsma, M. F. & Spek, C. A. The dual role of C/EBPδ in cancer. Crit. Rev. Oncol. Hematol. 185, 103983 (2023).

40. Li, C.-F. et al. HMDB and 5-AzadC Combination Reverses Tumor Suppressor CCAAT/Enhancer-Binding Protein Delta to Strengthen the Death of Liver Cancer Cells. Mol. Cancer Ther. 14, 2623–2633 (2015).

41. Yang, K. et al. FOXM1 promotes the growth and metastasis of colorectal cancer via activation of β-catenin signaling pathway. Cancer Manag. Res. 11, 3779–3790 (2019).

42. Zhang, X. et al. The tumor suppressive role of miRNA-370 by targeting FoxM1 in acute myeloid leukemia. Mol. Cancer 11, 56 (2012).

43. Liu, F. et al. Inhibitor of DNA binding 2 knockdown inhibits the growth and liver metastasis of colorectal cancer. Gene 819, 146240 (2022).

44. Kicheva, A. et al. Coordination of progenitor specification and growth in mouse and chick spinal cord. Science 345, 1254927 (2014).

45. Sagner, A. et al. Olig2 and Hes regulatory dynamics during motor neuron differentiation revealed by single cell transcriptomics. PLOS Biol. 16, e2003127 (2018).

46. Ericson, J. et al. Pax6 Controls Progenitor Cell Identity and Neuronal Fate in Response to Graded Shh Signaling. Cell 90, 169–180 (1997).

47. Usui, N. et al. Role of motoneuron-derived neurotrophin 3 in survival and axonal projection of sensory neurons during neural circuit formation. Development 139, 1125–1132 (2012).

48. Kim, M. et al. Motor neuron cell bodies are actively positioned by Slit/Robo repulsion and Netrin/DCC attraction. Dev. Biol. 399, 68–79 (2015).

49. Chédotal, A. Roles of axon guidance molecules in neuronal wiring in the developing spinal cord. Nat. Rev. Neurosci. 20, 380–396 (2019).

50. Qiu, X. et al. Mapping transcriptomic vector fields of single cells. Cell 185, 690-711.e45 (2022).

51. Virshup, I., Rybakov, S., Theis, F. J., Angerer, P. & Wolf, F. A. anndata: Access and store annotated datamatrices. J. Open Source Softw. 9, 4371 (2024).

52. Wolf, F. A., Angerer, P. & Theis, F. J. SCANPY: large-scale single-cell gene expression data analysis. Genome Biol. 19, 15 (2018).

53. Peyré, G. & Cuturi, M. Computational Optimal Transport. Preprint at 10.48550/arXiv.1803.00567 (2020).

54. Cuturi, M. Sinkhorn Distances: Lightspeed Computation of Optimal Transportation Distances. Preprint at 10.48550/arXiv.1306.0895 (2013).

55. Jordan, R., Kinderlehrer, D. & Otto, F. The Variational Formulation of the Fokker--Planck Equation. SIAM J. Math. Anal. 29, 1–17 (1998).

56. Cuturi, M. et al. Optimal Transport Tools (OTT): A JAX Toolbox for all things Wasserstein. Preprint at 10.48550/arXiv.2201.12324 (2022).

57. google/flax. Google (2026).

58. Gao, M., Qiao, C. & Huang, Y. UniTVelo: temporally unified RNA velocity reinforces single-cell trajectory inference. Nat. Commun. 13, 6586 (2022).

59. Pedregosa, F. et al. Scikit-learn: Machine Learning in Python. Mach. Learn. PYTHON.

