## Supplementary figures and tables for "Inferring Cell Fate Trajectories in Time-Resolved Metabolic RNA Labeling data"

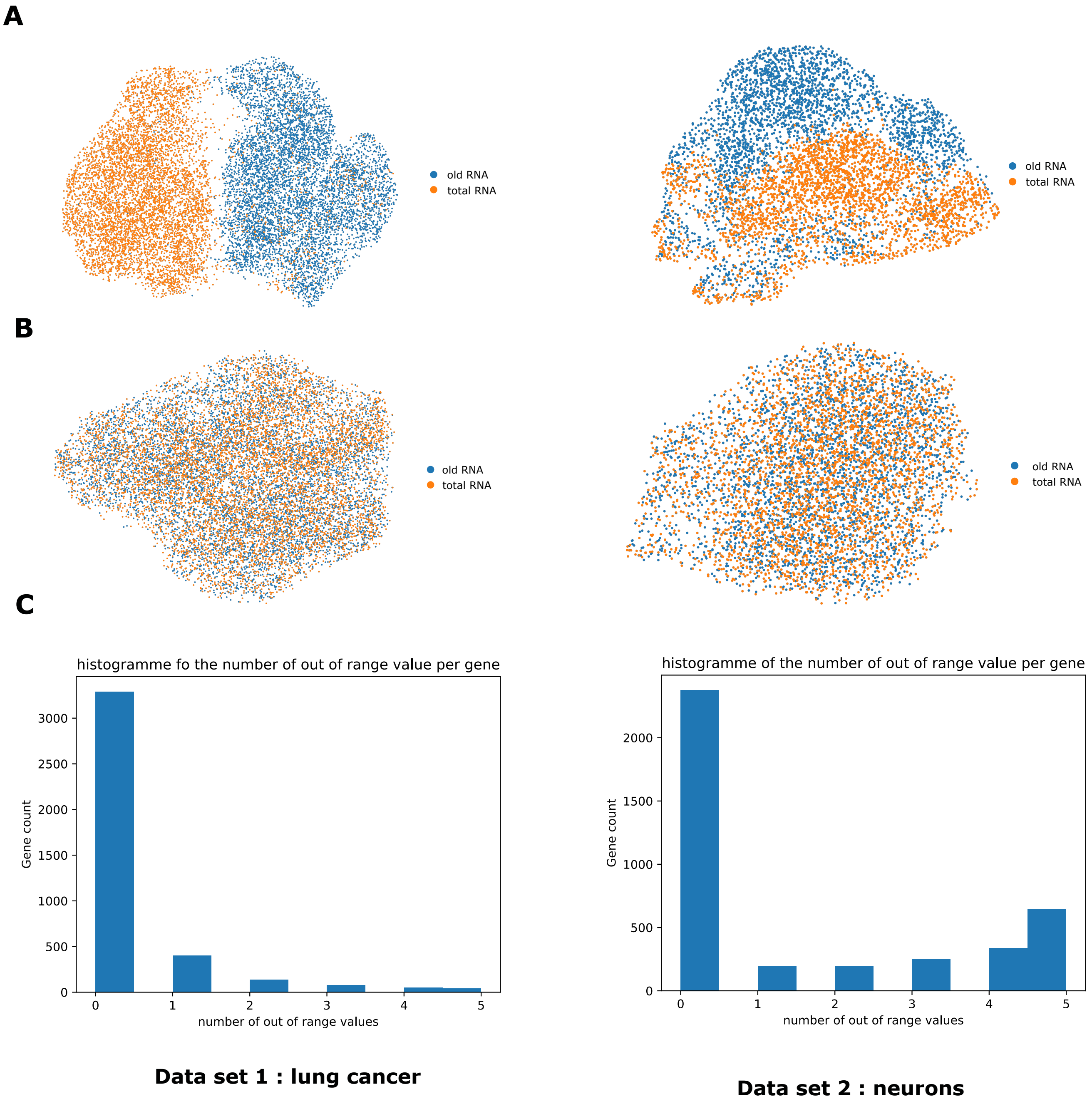


**Supplementary Figure 1 : A. Joint Umap before correction B. Joint Umap after correction C. Number of out of range value degradation estimated for each gene (one degradation rate per interval)**

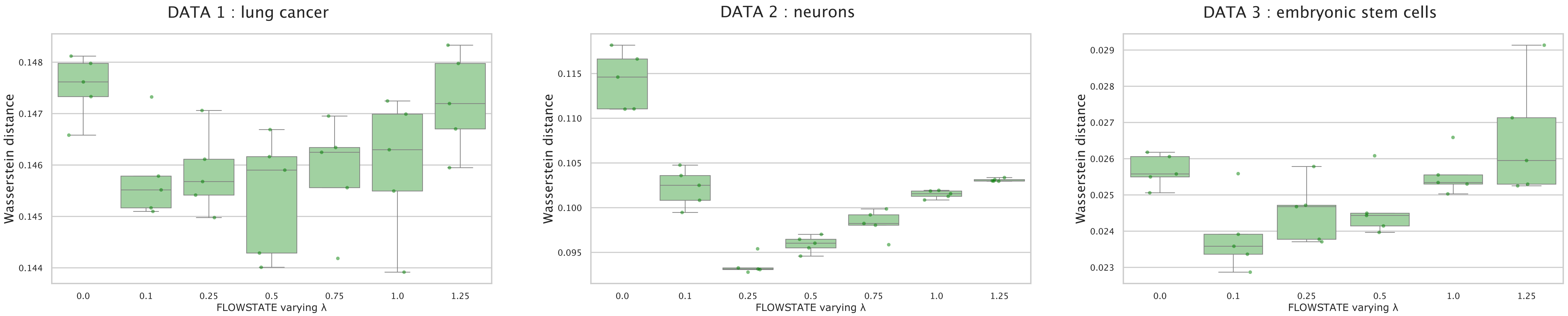


**Supplementary Figure 2. Wasserstein loss between prediction and ground truth at the last time point for varying value of λ for all three benchmarking datasets**

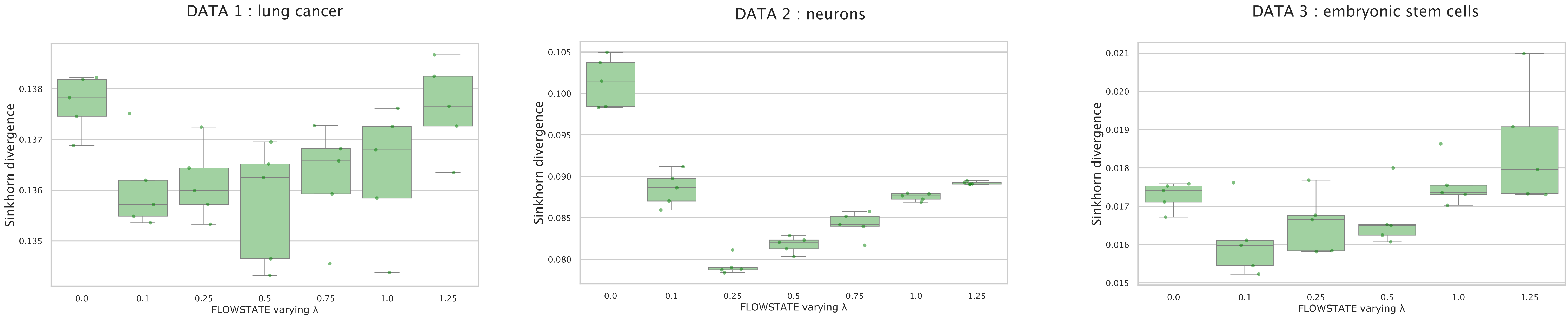


**Supplementary Figure 3. Sinkhorn loss between prediction and ground truth at the last time point for varying value of λ for all three benchmarking datasets**

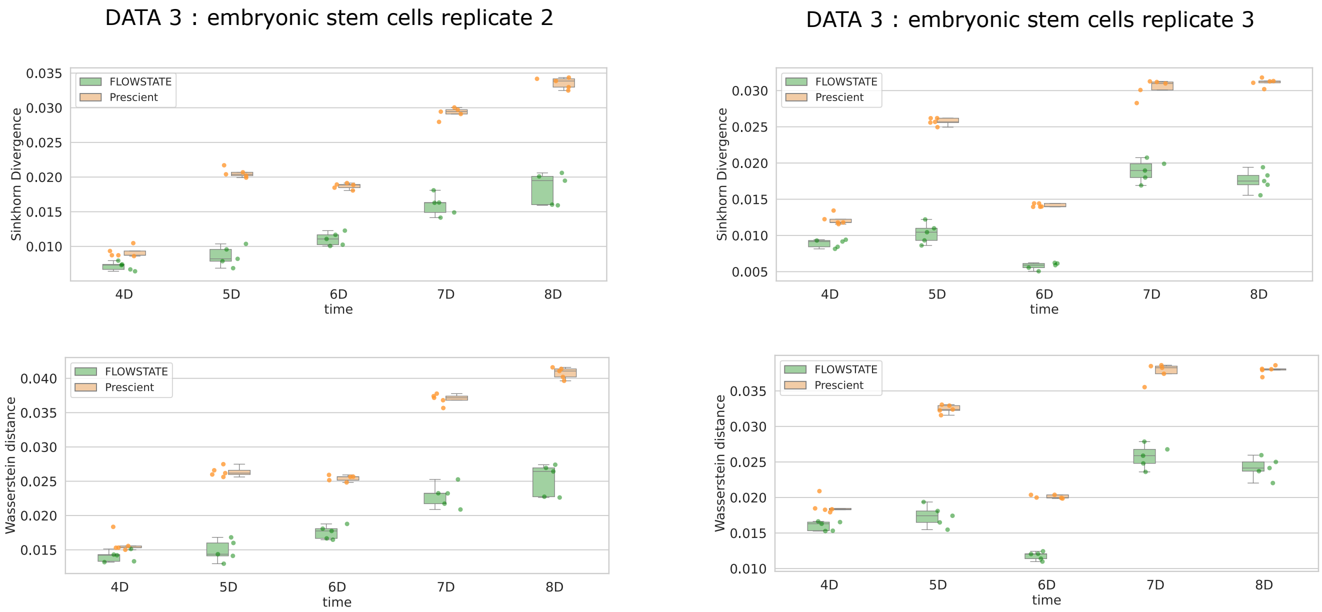


**Supplementary Figure 4. Comparaison of the sinkhorn divergence and the wasserstein loss between prediction and ground truth at every time point for replicates 2 and 3 of dataset 3 after training FLOWSTATE and Prescient on replicate 1**

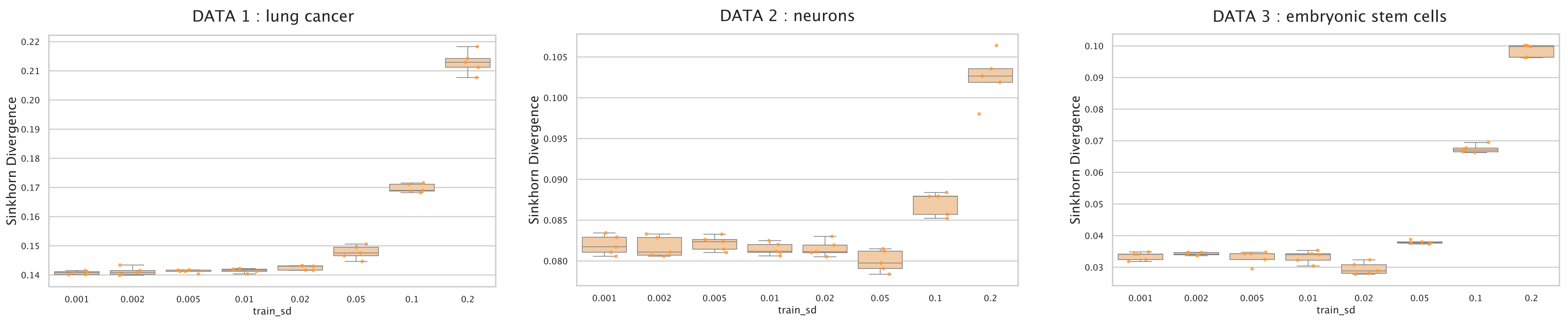


**Supplementary Figure 5. Prescient performances for the sinkhorn loss between prediction and ground truth at the last time point for varying value of train_sd, the noise parameter for all three benchmarking datasets**

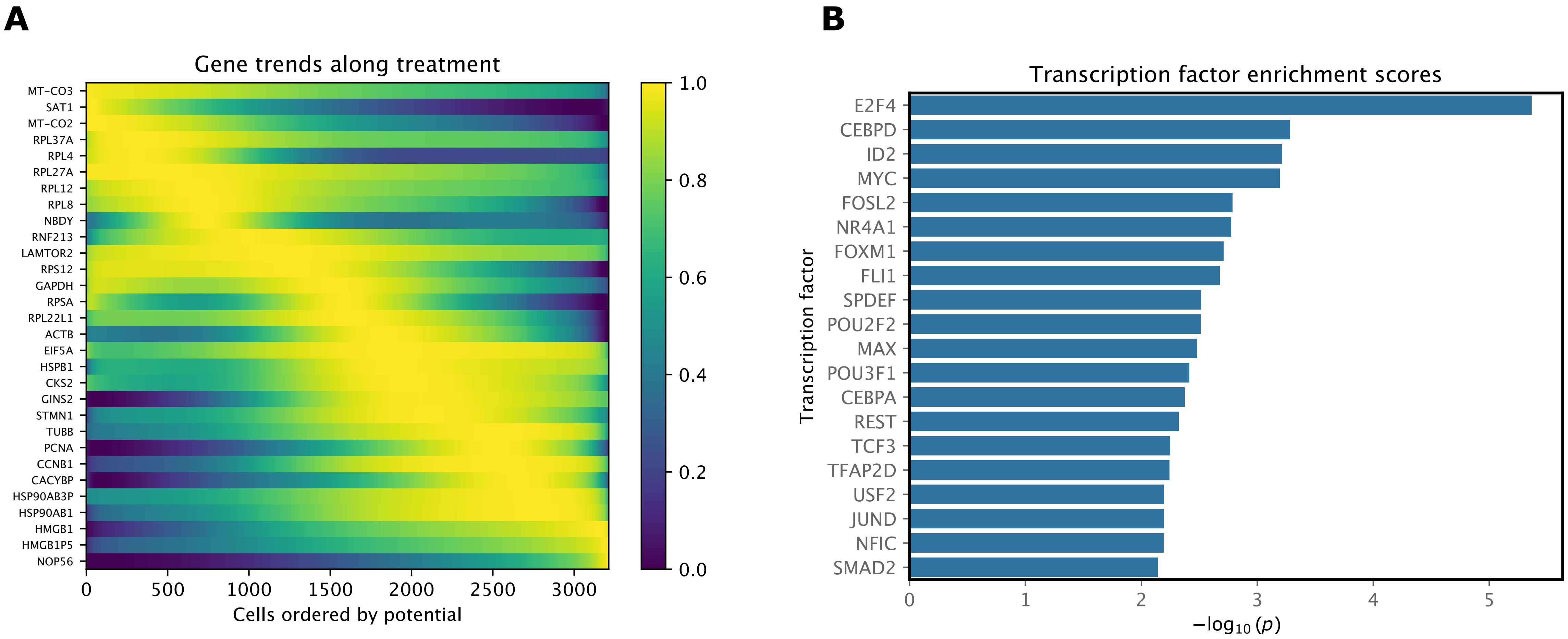


**Supplementary Figure 6. Trajectory inference on colorectal cancer cells. A. Normalized gene expression for the genes most correlated with the potential. Expression regressed using a spline model along the potential computed by FLOWSTATE B. Enrichment score of the top 20 transcription factors targeting the genes most correlated with the potential. A one-sided Wilcoxon rank-sum test is used to report P values.**

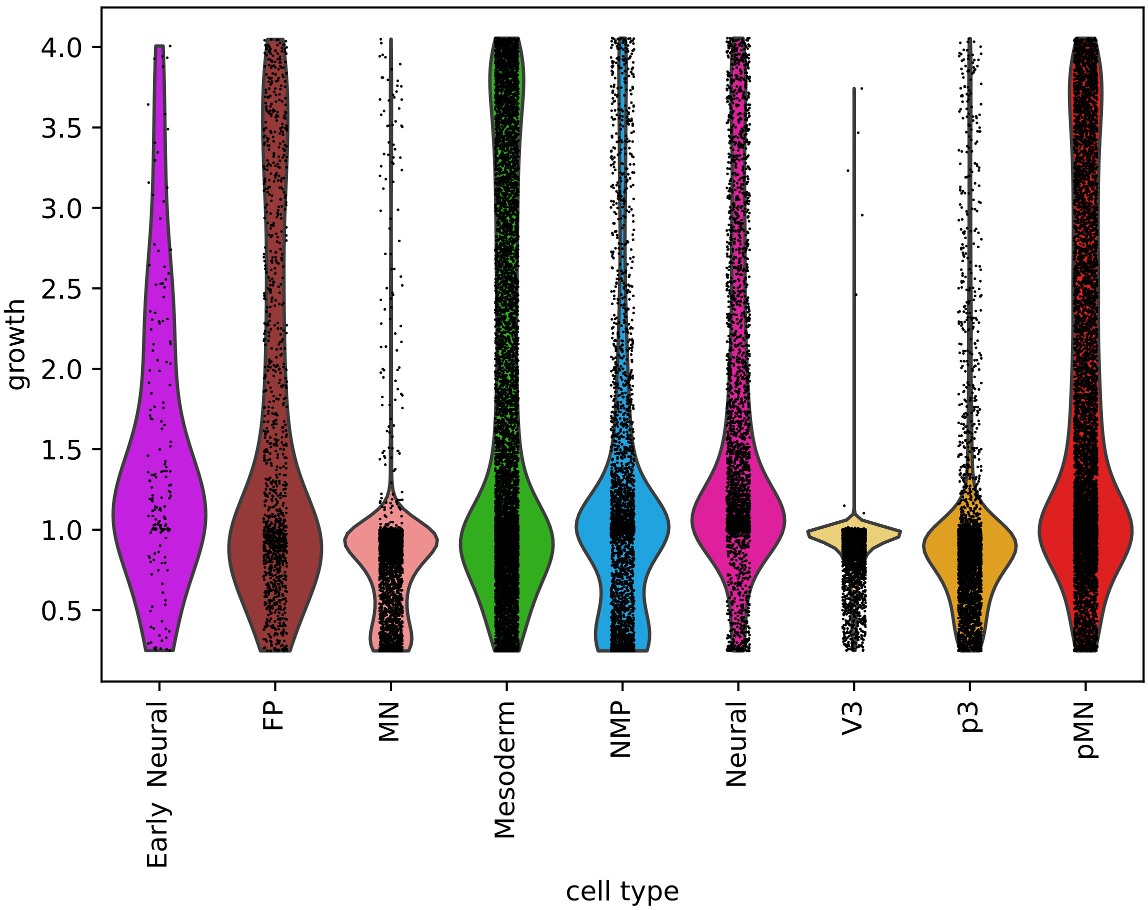


**Supplementary Figure 7. Growth rate per cell type for mouse embryonic stem cells**

### From 3D to 4D

| Source Type | Predicted Type | MN | Mesoderm | NMP | Neural | pMN |
|---|---|---|---|---|---|---|
| | Early_Neural | 0.000 | 0.000 | 0.000 | 0.000 | 0.000 |
| | FP | 0.000 | 0.000 | 0.000 | 0.000 | 0.000 |
| | MN | 0.000 | 0.000 | 0.000 | 0.000 | 0.000 |
| | Mesoderm | 0.000 | 0.000 | 0.000 | 0.000 | 0.000 |
| | NMP | **0.001** | **0.240** | **0.164** | **0.593** | **0.002** |
| | Neural | 0.000 | 0.000 | 0.000 | 0.000 | 0.000 |
| | V3 | 0.000 | 0.000 | 0.000 | 0.000 | 0.000 |
| | p3 | 0.000 | 0.000 | 0.000 | 0.000 | 0.000 |
| | pMN | 0.000 | 0.000 | 0.000 | 0.000 | 0.000 |

### From 4D to 5D

| Source Type | Predicted Type | MN | Mesoderm | Neural | V3 | p3 | pMN |
|---|---|---|---|---|---|---|---|
| | Early_Neural | 0.000 | 0.000 | 0.000 | 0.000 | 0.000 | 0.000 |
| | FP | 0.000 | 0.000 | 0.000 | 0.000 | 0.000 | 0.000 |
| | MN | 0.000 | 0.000 | 0.000 | 0.000 | 0.000 | 0.000 |
| | Mesoderm | **0.001** | **0.120** | **0.001** | 0.000 | 0.000 | **0.011** |
| | NMP | 0.000 | 0.000 | 0.000 | 0.000 | 0.000 | 0.000 |
| | Neural | **0.155** | **0.077** | **0.158** | 0.000 | **0.002** | **0.476** |
| | V3 | 0.000 | 0.000 | 0.000 | 0.000 | 0.000 | 0.000 |
| | p3 | 0.000 | 0.000 | 0.000 | 0.000 | 0.000 | 0.000 |
| | pMN | 0.000 | 0.000 | 0.000 | 0.000 | 0.000 | 0.000 |

### From 5D to 6D

| Source Type | Predicted Type | FP | MN | Mesoderm | V3 | p3 | pMN |
|---|---|---|---|---|---|---|---|
| | Early_Neural | 0.000 | 0.000 | 0.000 | 0.000 | 0.000 | 0.000 |
| | FP | 0.000 | 0.000 | 0.000 | 0.000 | 0.000 | 0.000 |
| | MN | 0.000 | **0.085** | 0.000 | **0.007** | 0.000 | **0.048** |
| | Mesoderm | 0.000 | **0.001** | **0.343** | 0.000 | 0.000 | **0.023** |
| | NMP | 0.000 | 0.000 | 0.000 | 0.000 | 0.000 | 0.000 |
| | Neural | 0.000 | 0.000 | 0.000 | 0.000 | 0.000 | 0.000 |
| | V3 | 0.000 | 0.000 | 0.000 | 0.000 | 0.000 | 0.000 |
| | p3 | 0.000 | 0.000 | 0.000 | 0.000 | 0.000 | 0.000 |
| | pMN | **0.016** | **0.014** | **0.001** | 0.000 | **0.029** | **0.433** |

### From 6D to 7D

| Source Type | Predicted Type | FP | MN | Mesoderm | Neural | V3 | p3 | pMN |
|---|---|---|---|---|---|---|---|---|
| | Early_Neural | 0.000 | 0.000 | 0.000 | 0.000 | 0.000 | 0.000 | 0.000 |
| | FP | **0.011** | 0.000 | 0.000 | 0.000 | 0.000 | **0.001** | 0.000 |
| | MN | 0.000 | **0.073** | 0.000 | 0.000 | **0.007** | 0.000 | 0.000 |
| | Mesoderm | 0.000 | 0.000 | **0.111** | 0.000 | 0.000 | 0.000 | **0.035** |
| | NMP | 0.000 | 0.000 | 0.000 | 0.000 | 0.000 | 0.000 | 0.000 |
| | Neural | 0.000 | 0.000 | 0.000 | 0.000 | 0.000 | 0.000 | 0.000 |
| | V3 | 0.000 | 0.000 | 0.000 | 0.000 | **0.001** | 0.000 | 0.000 |
| | p3 | 0.000 | 0.000 | 0.000 | 0.000 | **0.004** | **0.010** | 0.000 |
| | pMN | **0.012** | **0.052** | **0.001** | 0.000 | **0.011** | **0.106** | **0.565** |

### From 7D to 8D

| Source Type | Predicted Type | FP | MN | Mesoderm | V3 | p3 | pMN |
|---|---|---|---|---|---|---|---|
| | Early_Neural | 0.000 | 0.000 | 0.000 | 0.000 | 0.000 | 0.000 |
| | FP | **0.032** | 0.000 | 0.000 | 0.000 | **0.001** | 0.000 |
| | MN | 0.000 | **0.109** | 0.000 | **0.004** | 0.000 | **0.001** |
| | Mesoderm | 0.000 | 0.000 | **0.016** | 0.000 | **0.001** | **0.008** |
| | NMP | 0.000 | 0.000 | 0.000 | 0.000 | 0.000 | 0.000 |
| | Neural | 0.000 | 0.000 | 0.000 | 0.000 | 0.000 | 0.000 |
| | V3 | 0.000 | 0.000 | 0.000 | **0.051** | 0.000 | 0.000 |
| | p3 | **0.001** | 0.000 | 0.000 | **0.047** | **0.116** | **0.002** |
| | pMN | **0.018** | **0.042** | 0.000 | **0.029** | **0.170** | **0.352** |

**Supplementary Table 1. Transition probabilities between cell types across consecutive time points in embryonic stem cells (Dataset 3) after training FLOWSTATE with the last timepoint removed. Probabilities are normalized per time point transition.**

### From 3D to 4D

| | Predicted Type | MN | Mesoderm | Neural | pMN |
|---|---|---|---|---|---|
| Source Type | Early_Neural | 0.000 | 0.000 | 0.000 | 0.000 |
| | FP | 0.000 | 0.000 | 0.000 | 0.000 |
| | MN | 0.000 | 0.000 | 0.000 | 0.000 |
| | Mesoderm | 0.000 | 0.000 | 0.000 | 0.000 |
| | NMP | 0.000 | **0.318** | **0.675** | **0.007** |
| | Neural | 0.000 | 0.000 | 0.000 | 0.000 |
| | V3 | 0.000 | 0.000 | 0.000 | 0.000 |
| | p3 | 0.000 | 0.000 | 0.000 | 0.000 |
| | pMN | 0.000 | 0.000 | 0.000 | 0.000 |

### From 4D to 5D

| | Predicted Type | MN | Mesoderm | Neural | pMN |
|---|---|---|---|---|---|
| Source Type | Early_Neural | 0.000 | 0.000 | 0.000 | 0.000 |
| | FP | 0.000 | 0.000 | 0.000 | 0.000 |
| | MN | 0.000 | 0.000 | 0.000 | 0.000 |
| | Mesoderm | 0.000 | **0.119** | **0.008** | **0.006** |
| | NMP | 0.000 | 0.000 | 0.000 | 0.000 |
| | Neural | **0.013** | 0.000 | **0.737** | **0.117** |
| | V3 | 0.000 | 0.000 | 0.000 | 0.000 |
| | p3 | 0.000 | 0.000 | 0.000 | 0.000 |
| | pMN | 0.000 | 0.000 | 0.000 | 0.000 |

### From 5D to 6D

| | Predicted Type | FP | MN | Mesoderm | Neural | p3 | pMN |
|---|---|---|---|---|---|---|---|
| Source Type | Early_Neural | 0.000 | 0.000 | 0.000 | 0.000 | 0.000 | 0.000 |
| | FP | 0.000 | 0.000 | 0.000 | 0.000 | 0.000 | 0.000 |
| | MN | 0.000 | **0.117** | 0.000 | 0.000 | **0.002** | **0.020** |
| | Mesoderm | 0.000 | 0.000 | **0.364** | 0.000 | 0.000 | **0.003** |
| | NMP | 0.000 | 0.000 | 0.000 | 0.000 | 0.000 | 0.000 |
| | Neural | 0.000 | 0.000 | 0.000 | 0.000 | 0.000 | 0.000 |
| | V3 | 0.000 | 0.000 | 0.000 | 0.000 | 0.000 | 0.000 |
| | p3 | 0.000 | 0.000 | 0.000 | 0.000 | 0.000 | 0.000 |
| | pMN | **0.002** | **0.003** | 0.000 | 0.000 | **0.002** | **0.487** |

### From 6D to 7D

| | Predicted Type | FP | MN | Mesoderm | Neural | V3 | p3 | pMN |
|---|---|---|---|---|---|---|---|---|
| Source Type | Early_Neural | 0.000 | 0.000 | 0.000 | 0.000 | 0.000 | 0.000 | 0.000 |
| | FP | **0.010** | 0.000 | 0.000 | 0.000 | 0.000 | 0.000 | **0.002** |
| | MN | 0.000 | **0.079** | 0.000 | 0.000 | 0.000 | 0.000 | **0.002** |
| | Mesoderm | 0.000 | 0.000 | **0.145** | 0.000 | 0.000 | 0.000 | 0.000 |
| | NMP | 0.000 | 0.000 | 0.000 | 0.000 | 0.000 | 0.000 | 0.000 |
| | Neural | 0.000 | 0.000 | 0.000 | 0.000 | 0.000 | 0.000 | 0.000 |
| | V3 | 0.000 | 0.000 | 0.000 | 0.000 | 0.000 | 0.000 | 0.000 |
| | p3 | 0.000 | 0.000 | 0.000 | 0.000 | **0.001** | **0.012** | **0.002** |
| | pMN | **0.001** | **0.013** | 0.000 | 0.000 | 0.000 | **0.018** | **0.714** |

### From 7D to 8D

| | Predicted Type | FP | MN | Mesoderm | V3 | p3 | pMN |
|---|---|---|---|---|---|---|---|
| Source Type | Early_Neural | 0.000 | 0.000 | 0.000 | 0.000 | 0.000 | 0.000 |
| | FP | **0.030** | 0.000 | 0.000 | 0.000 | **0.001** | **0.002** |
| | MN | 0.000 | **0.113** | 0.000 | 0.000 | **0.001** | **0.001** |
| | Mesoderm | 0.000 | 0.000 | **0.025** | 0.000 | 0.000 | 0.000 |
| | NMP | 0.000 | 0.000 | 0.000 | 0.000 | 0.000 | 0.000 |
| | Neural | 0.000 | 0.000 | 0.000 | 0.000 | 0.000 | 0.000 |
| | V3 | 0.000 | **0.002** | 0.000 | **0.049** | 0.000 | 0.000 |
| | p3 | **0.001** | 0.000 | 0.000 | **0.005** | **0.155** | **0.005** |
| | pMN | **0.013** | **0.007** | 0.000 | 0.000 | **0.069** | **0.522** |

**Supplementary Table 2. Transition probabilities between cell types across consecutive time points in embryonic stem cells (Dataset 3) after training Prescient with the last timepoint removed. Probabilities are normalized per time point transition.**

| Gene_set | Term | Overlap | P-value | Adjusted P-value | Odds Ratio | Combined Score | Genes |
|---|---|---|---|---|---|---|---|
| GO_Biological_Process_2025 | Cytoplasmic Translation (GO:0002181) | 7/101 | 0.000000 | 0.000003 | 34.386937 | 652.935082 | RPS15;RPL12;RPSA;RPL22L1;RPS2;RPL8;RPS12 |
| GO_Biological_Process_2025 | Translation (GO:0006412) | 7/232 | 0.000002 | 0.000400 | 14.271318 | 189.703069 | RPS15;RPL12;RPSA;RPL22L1;RPS2;RPL8;RPS12 |
| GO_Biological_Process_2025 | Regulation of Mitotic Sister Chromatid Separation (GO:0010965) | 3/12 | 0.000003 | 0.000503 | 141.425532 | 1790.106623 | CCNB1;CDCA8;AURKB |
| GO_Biological_Process_2025 | Gene Expression (GO:0010467) | 8/381 | 0.000004 | 0.000510 | 9.997191 | 123.532619 | RPS15;RPL12;HNRNPD;RPSA;RPS2;RPL8;PTMA;RPS12 |
| GO_Biological_Process_2025 | Macromolecule Biosynthetic Process (GO:0009059) | 6/189 | 0.000007 | 0.000701 | 14.729508 | 174.018535 | RPS15;RPL12;RPSA;RPS2;RPL8;RPS12 |
| GO_Biological_Process_2025 | Regulation of Mitotic Cell Cycle Spindle Assembly Checkpoint (GO:0090266) | 3/20 | 0.000016 | 0.001285 | 74.842303 | 825.250012 | CCNB1;CDCA8;AURKB |
| GO_Biological_Process_2025 | Ribosomal Small Subunit Biogenesis (GO:0042274) | 4/85 | 0.000060 | 0.004082 | 21.330113 | 207.254492 | NOP56;RPS15;RPSA;RPS12 |
| GO_Biological_Process_2025 | Mitotic Spindle Organization (GO:0007052) | 4/90 | 0.000075 | 0.004466 | 20.084934 | 190.667047 | CCNB1;STMN1;CDCA8;AURKB |
| GO_Biological_Process_2025 | Positive Regulation of Leukocyte Differentiation (GO:1902107) | 2/7 | 0.000128 | 0.006048 | 166.208333 | 1490.324070 | LGALS1;HMGB1 |
| GO_Biological_Process_2025 | Regulation of Mitotic Cytokinesis (GO:1902412) | 2/7 | 0.000128 | 0.006048 | 166.208333 | 1490.324070 | CDCA8;AURKB |
| **Gene_set** | **Term** | **Overlap** | **P-value** | **Adjusted P-value** | **Odds Ratio** | **Combined Score** | **Genes** |
| GO_Cellular_Component_2025 | Cytosolic Small Ribosomal Subunit (GO:0022627) | 4/40 | 0.000003 | 0.000141 | 48.101449 | 612.423721 | RPS15;RPSA;RPS2;RPS12 |
| GO_Cellular_Component_2025 | Small Ribosomal Subunit (GO:0015935) | 4/41 | 0.000003 | 0.000141 | 46.799060 | 591.123728 | RPS15;RPSA;RPS2;RPS12 |
| **Gene_set** | **Term** | **Overlap** | **P-value** | **Adjusted P-value** | **Odds Ratio** | **Combined Score** | **Genes** |
| GO_Molecular_Function_2025 | DNA Polymerase Binding (GO:0070182) | 3/20 | 0.000016 | 0.001382 | 74.842303 | 825.250012 | HSP90AB1;PCNA;HMGB1 |
| **Gene_set** | **Term** | **Overlap** | **P-value** | **Adjusted P-value** | **Odds Ratio** | **Combined Score** | **Genes** |
| MSigDB_Hallmark_2020 | E2F Targets | 11/200 | 0.000000 | 0.000000 | 29.490028 | 794.169656 | NOP56;DUT;PCNA;TUBB;UBE2T;STMN1;HNRNPD;CKS2;CDCA8;SSRP1;AURKB |
| MSigDB_Hallmark_2020 | Myc Targets V1 | 7/200 | 0.000001 | 0.000007 | 16.664538 | 238.067958 | NOP56;DUT;PCNA;HSP90AB1;HNRNPD;RPS2;PSMA7 |
| MSigDB_Hallmark_2020 | G2-M Checkpoint | 5/200 | 0.000140 | 0.000979 | 11.256410 | 99.901053 | GINS2;STMN1;HNRNPD;CKS2;AURKB |
| MSigDB_Hallmark_2020 | DNA Repair | 4/150 | 0.000535 | 0.002811 | 11.795116 | 88.847050 | DUT;PCNA;POLR3GL;SSRP1 |
| MSigDB_Hallmark_2020 | p53 Pathway | 4/200 | 0.001560 | 0.006551 | 8.763975 | 56.642851 | LDHB;PCNA;SAT1;RPS12 |
| KEGG_2021_Human | Ribosome | 7/158 | 0.000000 | 0.000012 | 21.344987 | 339.071541 | RPS15;RPL12;RPSA;RPL22L1;RPS2;RPL8;RPS12 |
| KEGG_2021_Human | Coronavirus disease | 7/232 | 0.000002 | 0.000078 | 14.271318 | 189.703069 | RPS15;RPL12;RPSA;RPL22L1;RPS2;RPL8;RPS12 |
| **Gene_set** | **Term** | **Overlap** | **P-value** | **Adjusted P-value** | **Odds Ratio** | **Combined Score** | **Genes** |
| WikiPathways_2024_Human | Cytoplasmic Ribosomal Proteins WP477 | 6/88 | 0.000000 | 0.000009 | 33.039911 | 538.682843 | RPS15;RPL12;RPSA;RPS2;RPL8;RPS12 |
| WikiPathways_2024_Human | Retinoblastoma Gene In Cancer WP2446 | 4/87 | 0.000066 | 0.003400 | 20.814039 | 200.345485 | CCNB1;PCNA;STMN1;HMGB1 |
| **Gene_set** | **Term** | **Overlap** | **P-value** | **Adjusted P-value** | **Odds Ratio** | **Combined Score** | **Genes** |
| Reactome_Pathways_2024 | rRNA Processing | 13/237 | 0.000000 | 0.000000 | 30.940878 | 983.431656 | NOP56;MT-ND4;MT-CO1;RPL12;RPSA;RPL8;RPS15;RPL22L1;MT-CO2;RPS2;MT-CO3;MT-CYB;RPS12 |
| Reactome_Pathways_2024 | Metabolism of RNA | 17/761 | 0.000000 | 0.000000 | 13.298387 | 359.133571 | NOP56;MT-ND4;MT-CO1;RPL12;RPSA;RPL8;ACTB;PSMA7;RPS15;HNRNPD;RPL22L1;MT-CO2;SNRPF;RPS2;MT-CO3;MT-CYB;RPS12 |
| Reactome_Pathways_2024 | FASTK Family Proteins Regulate Processing and Stability of Mitochondrial RNAs | 5/19 | 0.000000 | 0.000000 | 158.222222 | 3295.518911 | MT-ND4;MT-CO1;MT-CO2;MT-CO3;MT-CYB |
| Reactome_Pathways_2024 | Peptide Chain Elongation | 7/94 | 0.000000 | 0.000000 | 37.166800 | 724.517603 | RPS15;RPL12;RPSA;RPL22L1;RPS2;RPL8;RPS12 |
| Reactome_Pathways_2024 | Mitochondrial RNA Degradation | 5/25 | 0.000000 | 0.000000 | 110.722222 | 2139.189535 | MT-ND4;MT-CO1;MT-CO2;MT-CO3;MT-CYB |
| Reactome_Pathways_2024 | Eukaryotic Translation Termination | 7/98 | 0.000000 | 0.000000 | 35.525939 | 682.096418 | RPS15;RPL12;RPSA;RPL22L1;RPS2;RPL8;RPS12 |
| Reactome_Pathways_2024 | Selenocysteine Synthesis | 7/98 | 0.000000 | 0.000000 | 35.525939 | 682.096418 | RPS15;RPL12;RPSA;RPL22L1;RPS2;RPL8;RPS12 |
| Reactome_Pathways_2024 | Eukaryotic Translation Elongation | 7/99 | 0.000000 | 0.000000 | 35.138018 | 672.137865 | RPS15;RPL12;RPSA;RPL22L1;RPS2;RPL8;RPS12 |
| Reactome_Pathways_2024 | Nonsense Mediated Decay (NMD) Independent of the Exon Junction Complex (EJC) | 7/100 | 0.000000 | 0.000000 | 34.758440 | 662.420162 | RPS15;RPL12;RPSA;RPL22L1;RPS2;RPL8;RPS12 |
| Reactome_Pathways_2024 | Cellular Response to Starvation | 8/161 | 0.000000 | 0.000000 | 24.646125 | 466.730495 | RPS15;RPL12;RPSA;RPL22L1;LAMTOR2;RPS2;RPL8;RPS12 |
| Reactome_Pathways_2024 | Viral mRNA Translation | 7/102 | 0.000000 | 0.000000 | 34.023256 | 643.674760 | RPS15;RPL12;RPSA;RPL22L1;RPS2;RPL8;RPS12 |
| Reactome_Pathways_2024 | Regulation of Expression of SLITs and ROBOs | 8/164 | 0.000000 | 0.000000 | 24.168498 | 454.172499 | RPS15;RPL12;RPSA;RPL22L1;RPS2;RPL8;PSMA7;RPS12 |
| Reactome_Pathways_2024 | Response of EIF2AK4 (GCN2) to Amino Acid Deficiency | 7/106 | 0.000000 | 0.000000 | 32.642001 | 608.735381 | RPS15;RPL12;RPSA;RPL22L1;RPS2;RPL8;RPS12 |
| Reactome_Pathways_2024 | Formation of a Pool of Free 40S Subunits | 7/106 | 0.000000 | 0.000000 | 32.642001 | 608.735381 | RPS15;RPL12;RPSA;RPL22L1;RPS2;RPL8;RPS12 |
| Reactome_Pathways_2024 | L13a-mediated Translational Silencing of Ceruloplasmin Expression | 7/116 | 0.000000 | 0.000000 | 29.632387 | 533.941701 | RPS15;RPL12;RPSA;RPL22L1;RPS2;RPL8;RPS12 |
| Reactome_Pathways_2024 | GTP Hydrolysis and Joining of the 60S Ribosomal Subunit | 7/117 | 0.000000 | 0.000000 | 29.361522 | 527.304964 | RPS15;RPL12;RPSA;RPL22L1;RPS2;RPL8;RPS12 |
| Reactome_Pathways_2024 | Metabolism of Amino Acids and Derivatives | 10/358 | 0.000000 | 0.000000 | 14.081897 | 252.608248 | RPS15;SLC44A1;RPL12;RPSA;RPL22L1;RPS2;RPL8;SAT1;PSMA7;RPS12 |
| Reactome_Pathways_2024 | SRP-dependent Cotranslational Protein Targeting to Membrane | 7/119 | 0.000000 | 0.000000 | 28.834302 | 514.433921 | RPS15;RPL12;RPSA;RPL22L1;RPS2;RPL8;RPS12 |
| Reactome_Pathways_2024 | Nonsense Mediated Decay (NMD) Enhanced by the Exon Junction Complex (EJC) | 7/120 | 0.000000 | 0.000000 | 28.577691 | 508.191961 | RPS15;RPL12;RPSA;RPL22L1;RPS2;RPL8;RPS12 |
| Reactome_Pathways_2024 | Nonsense-Mediated Decay (NMD) | 7/120 | 0.000000 | 0.000000 | 28.577691 | 508.191961 | RPS15;RPL12;RPSA;RPL22L1;RPS2;RPL8;RPS12 |
| Reactome_Pathways_2024 | Major Pathway of rRNA Processing in the Nucleolus and Cytosol | 8/189 | 0.000000 | 0.000000 | 20.803999 | 367.825144 | NOP56;RPS15;RPL12;RPSA;RPL22L1;RPS2;RPL8;RPS12 |
| Reactome_Pathways_2024 | Selenoamino Acid Metabolism | 7/122 | 0.000000 | 0.000000 | 28.077856 | 496.077077 | RPS15;RPL12;RPSA;RPL22L1;RPS2;RPL8;RPS12 |
| Reactome_Pathways_2024 | Cap-dependent Translation Initiation | 7/124 | 0.000000 | 0.000001 | 27.595110 | 484.431637 | RPS15;RPL12;RPSA;RPL22L1;RPS2;RPL8;RPS12 |
| Reactome_Pathways_2024 | Eukaryotic Translation Initiation | 7/124 | 0.000000 | 0.000001 | 27.595110 | 484.431637 | RPS15;RPL12;RPSA;RPL22L1;RPS2;RPL8;RPS12 |
| Reactome_Pathways_2024 | rRNA Processing in the Nucleus and Cytosol | 8/199 | 0.000000 | 0.000001 | 19.704812 | 340.482482 | NOP56;RPS15;RPL12;RPSA;RPL22L1;RPS2;RPL8;RPS12 |
| Reactome_Pathways_2024 | rRNA Processing in the Mitochondrion | 5/38 | 0.000000 | 0.000001 | 67.060606 | 1146.665463 | MT-ND4;MT-CO1;MT-CO2;MT-CO3;MT-CYB |
| Reactome_Pathways_2024 | Cellular Responses to Stress | 13/787 | 0.000000 | 0.000001 | 8.704798 | 147.201614 | HSP90AB1;MT-CO1;RPL12;RPSA;RPL8;PSMA7;RPS15;RPL22L1;MT-CO2;LAMTOR2;RPS2;MT-CO3;RPS12 |
| Reactome_Pathways_2024 | Signaling by ROBO Receptors | 8/210 | 0.000000 | 0.000001 | 18.621405 | 313.990681 | RPS15;RPL12;RPSA;RPL22L1;RPS2;RPL8;PSMA7;RPS12 |
| Reactome_Pathways_2024 | tRNA Processing in the Mitochondrion | 5/42 | 0.000000 | 0.000001 | 59.798799 | 991.398491 | MT-ND4;MT-CO1;MT-CO2;MT-CO3;MT-CYB |
| Reactome_Pathways_2024 | Influenza Viral RNA Transcription and Replication | 7/149 | 0.000000 | 0.000001 | 22.708156 | 369.850548 | RPS15;RPL12;RPSA;RPL22L1;RPS2;RPL8;RPS12 |
| Reactome_Pathways_2024 | Cellular Responses to Stimuli | 13/887 | 0.000000 | 0.000003 | 7.668625 | 119.028432 | HSP90AB1;MT-CO1;RPL12;RPSA;RPL8;PSMA7;RPS15;RPL22L1;MT-CO2;LAMTOR2;RPS2;MT-CO3;RPS12 |
| Reactome_Pathways_2024 | Influenza Infection | 7/169 | 0.000000 | 0.000003 | 19.884582 | 306.740459 | RPS15;RPL12;RPSA;RPL22L1;RPS2;RPL8;RPS12 |
| Reactome_Pathways_2024 | Metabolism of Proteins | 19/2067 | 0.000000 | 0.000003 | 5.357516 | 82.051381 | EIF5A;NOP56;PCNA;MT-CO1;RPL12;CDCA8;RPSA;RPL8;AURKB;ACTB;PSMA7;RPS15;LGALS1;UBE2T;RPL22L1;MT-CO2;RPS2;PFDN5;RPS12 |
| Reactome_Pathways_2024 | SARS-CoV-2 Modulates Host Translation Machinery | 5/55 | 0.000000 | 0.000004 | 44.222222 | 671.950687 | RPS15;RPSA;SNRPF;RPS2;RPS12 |
| Reactome_Pathways_2024 | Metabolism | 19/2181 | 0.000001 | 0.000008 | 5.042702 | 72.996434 | DUT;MT-ND4;HSP90AB1;SLC44A1;MT-CO1;RPL12;RPSA;RPL8;SAT1;PSMA7;RPS15;LDHB;RPL22L1;MT-CO2;RPS2;MT-CO3;MT-CYB;GAPDH;RPS12 |
| Reactome_Pathways_2024 | Axon Guidance | 10/541 | 0.000001 | 0.000011 | 9.142655 | 128.947928 | RPS15;HSP90AB1;RPL12;RPSA;RPL22L1;RPS2;RPL8;ACTB;PSMA7;RPS12 |
| Reactome_Pathways_2024 | Viral Infection Pathways | 13/1029 | 0.000001 | 0.000014 | 6.547723 | 90.566630 | DUT;HSP90AB1;TUBB;RPL12;RPSA;SSRP1;RPL8;PSMA7;RPS15;RPL22L1;SNRPF;RPS2;RPS12 |
| Reactome_Pathways_2024 | Nervous System Development | 10/567 | 0.000001 | 0.000016 | 8.704219 | 119.063506 | RPS15;HSP90AB1;RPL12;RPSA;RPL22L1;RPS2;RPL8;ACTB;PSMA7;RPS12 |
| Reactome_Pathways_2024 | Infectious Disease | 14/1311 | 0.000003 | 0.000033 | 5.592864 | 72.069684 | DUT;HSP90AB1;TUBB;RPL12;RPSA;SSRP1;RPL8;ACTB;PSMA7;RPS15;RPL22L1;SNRPF;RPS2;RPS12 |
| Reactome_Pathways_2024 | Transcriptional Regulation by TP53 | 8/362 | 0.000003 | 0.000038 | 10.543987 | 134.268367 | CCNB1;PCNA;MT-CO1;MT-CO2;LAMTOR2;SSRP1;MT-CO3;AURKB |
| Reactome_Pathways_2024 | SARS-CoV-1 Modulates Host Translation Machinery | 4/41 | 0.000003 | 0.000041 | 46.799060 | 591.123728 | RPS15;RPSA;RPS2;RPS12 |
| Reactome_Pathways_2024 | Cell Cycle, Mitotic | 9/516 | 0.000005 | 0.000056 | 8.418098 | 103.424210 | GINS2;CCNB1;PCNA;HSP90AB1;TUBB;CDCA8;CENPN;AURKB;PSMA7 |
| Reactome_Pathways_2024 | Formation of the Ternary Complex, and Subsequently, the 43S Complex | 4/53 | 0.000009 | 0.000111 | 35.316770 | 409.369986 | RPS15;RPSA;RPS2;RPS12 |
| Reactome_Pathways_2024 | Translation | 7/302 | 0.000010 | 0.000112 | 10.846275 | 125.358245 | RPS15;RPL12;RPSA;RPL22L1;RPS2;RPL8;RPS12 |
| Reactome_Pathways_2024 | Ribosomal Scanning and Start Codon Recognition | 4/60 | 0.000015 | 0.000170 | 30.891304 | 342.715837 | RPS15;RPSA;RPS2;RPS12 |
| Reactome_Pathways_2024 | Translation Initiation Complex Formation | 4/60 | 0.000015 | 0.000170 | 30.891304 | 342.715837 | RPS15;RPSA;RPS2;RPS12 |
| Reactome_Pathways_2024 | mRNA Activation Upon Binding of the Cap-Binding Complex and eIFs, and Subsequent Binding to 43S | 4/61 | 0.000016 | 0.000178 | 30.347826 | 334.683853 | RPS15;RPSA;RPS2;RPS12 |
| Reactome_Pathways_2024 | SARS-CoV-2-host Interactions | 6/218 | 0.000017 | 0.000179 | 12.695969 | 139.678642 | RPS15;HSP90AB1;RPSA;SNRPF;RPS2;RPS12 |
| Reactome_Pathways_2024 | Cell Cycle | 9/649 | 0.000029 | 0.000302 | 6.623095 | 69.256200 | GINS2;CCNB1;PCNA;HSP90AB1;TUBB;CDCA8;CENPN;AURKB;PSMA7 |
| Reactome_Pathways_2024 | tRNA Processing | 5/146 | 0.000031 | 0.000324 | 15.609929 | 161.817020 | MT-ND4;MT-CO1;MT-CO2;MT-CO3;MT-CYB |
| Reactome_Pathways_2024 | Disease | 16/2131 | 0.000037 | 0.000376 | 3.968294 | 40.457102 | DUT;HSP90AB1;TUBB;RPL12;RPSA;HMGB1;SSRP1;RPL8;ACTB;PSMA7;RPS15;CCNB1;RPL22L1;SNRPF;RPS2;RPS12 |
| Reactome_Pathways_2024 | Aerobic Respiration and Respiratory Electron Transport | 6/260 | 0.000045 | 0.000443 | 10.574087 | 105.875456 | LDHB;MT-ND4;MT-CO1;MT-CO2;MT-CO3;MT-CYB |
| Reactome_Pathways_2024 | Respiratory Electron Transport | 5/158 | 0.000046 | 0.000445 | 14.376906 | 143.604003 | MT-ND4;MT-CO1;MT-CO2;MT-CO3;MT-CYB |
| Reactome_Pathways_2024 | TP53 Regulates Metabolic Genes | 4/86 | 0.000063 | 0.000601 | 21.068929 | 203.751908 | MT-CO1;MT-CO2;LAMTOR2;MT-CO3 |
| Reactome_Pathways_2024 | SARS-CoV-2 Infection | 6/313 | 0.000125 | 0.001165 | 8.725052 | 78.436495 | RPS15;HSP90AB1;RPSA;SNRPF;RPS2;RPS12 |
| Reactome_Pathways_2024 | Mitotic Prometaphase | 5/203 | 0.000150 | 0.001376 | 11.084175 | 97.601253 | CCNB1;TUBB;CDCA8;CENPN;AURKB |
| Reactome_Pathways_2024 | SARS-CoV-1-host Interactions | 4/112 | 0.000176 | 0.001587 | 15.975845 | 138.115084 | RPS15;RPSA;RPS2;RPS12 |
| Reactome_Pathways_2024 | SUMOylation of DNA Replication Proteins | 3/46 | 0.000207 | 0.001833 | 29.550223 | 250.683702 | PCNA;CDCA8;AURKB |
| Reactome_Pathways_2024 | Mitotic Anaphase | 5/221 | 0.000223 | 0.001915 | 10.151235 | 85.374885 | CCNB1;CDCA8;CENPN;AURKB;PSMA7 |
| Reactome_Pathways_2024 | SARS-CoV Infections | 7/498 | 0.000225 | 0.001915 | 6.451617 | 54.191611 | RPS15;HSP90AB1;TUBB;RPSA;SNRPF;RPS2;RPS12 |
| Reactome_Pathways_2024 | Mitotic Metaphase and Anaphase | 5/222 | 0.000227 | 0.001915 | 10.103943 | 84.765886 | CCNB1;CDCA8;CENPN;AURKB;PSMA7 |
| Reactome_Pathways_2024 | Complex IV Assembly | 3/48 | 0.000235 | 0.001947 | 28.234043 | 235.932951 | MT-CO1;MT-CO2;MT-CO3 |
| Reactome_Pathways_2024 | Resolution of Sister Chromatid Cohesion | 4/126 | 0.000276 | 0.002255 | 14.132573 | 115.798260 | CCNB1;CDCA8;CENPN;AURKB |
| Reactome_Pathways_2024 | M Phase | 6/372 | 0.000317 | 0.002544 | 7.296572 | 58.791390 | CCNB1;TUBB;CDCA8;CENPN;AURKB;PSMA7 |
| Reactome_Pathways_2024 | RHO GTPases Activate Formins | 4/139 | 0.000402 | 0.003175 | 12.763285 | 99.811947 | CDCA8;CENPN;AURKB;ACTB |
| Reactome_Pathways_2024 | Cytoprotection by HMOX1 | 3/59 | 0.000433 | 0.003373 | 22.675532 | 175.613224 | MT-CO1;MT-CO2;MT-CO3 |
| Reactome_Pathways_2024 | Cell Cycle Checkpoints | 5/261 | 0.000478 | 0.003666 | 8.547743 | 65.358728 | CCNB1;CDCA8;CENPN;AURKB;PSMA7 |
| Reactome_Pathways_2024 | Gene Expression (Transcription) | 12/1615 | 0.000497 | 0.003756 | 3.614342 | 27.494571 | CCNB1;PCNA;POLR3GL;MT-CO1;MT-CO2;LAMTOR2;SNRPF;SSRP1;MT-CO3;AURKB;ACTB;PSMA7 |
| Reactome_Pathways_2024 | Developmental Biology | 11/1385 | 0.000512 | 0.003811 | 3.813235 | 28.896594 | RPS15;CCNB1;HSP90AB1;RPL12;RPSA;RPL22L1;RPS2;RPL8;ACTB;PSMA7;RPS12 |
| Reactome_Pathways_2024 | The Role of GTSE1 in G2 M Progression After G2 Checkpoint | 3/65 | 0.000576 | 0.004228 | 20.474949 | 152.738497 | CCNB1;HSP90AB1;PSMA7 |
| Reactome_Pathways_2024 | SARS-CoV-1 Infection | 4/157 | 0.000636 | 0.004601 | 11.251492 | 82.823300 | RPS15;RPSA;RPS2;RPS12 |
| Reactome_Pathways_2024 | APC C-mediated Degradation of Cell Cycle Proteins | 3/75 | 0.000875 | 0.006158 | 17.622340 | 124.091470 | CCNB1;AURKB;PSMA7 |
| Reactome_Pathways_2024 | Regulation of Mitotic Cell Cycle | 3/75 | 0.000875 | 0.006158 | 17.622340 | 124.091470 | CCNB1;AURKB;PSMA7 |
| Reactome_Pathways_2024 | TP53 Regulates Transcription of Genes Involved in G2 Cell Cycle Arrest | 2/18 | 0.000913 | 0.006345 | 51.911458 | 363.289855 | CCNB1;PCNA |
| Reactome_Pathways_2024 | Separation of Sister Chromatids | 4/177 | 0.000994 | 0.006813 | 9.940689 | 68.726994 | CDCA8;CENPN;AURKB;PSMA7 |
| Reactome_Pathways_2024 | G2 M Transition | 4/184 | 0.001148 | 0.007762 | 9.550725 | 64.658976 | CCNB1;HSP90AB1;TUBB;PSMA7 |
| Reactome_Pathways_2024 | Mitotic G2-G2 M Phases | 4/186 | 0.001194 | 0.007973 | 9.444816 | 63.564733 | CCNB1;HSP90AB1;TUBB;PSMA7 |

**Supplementary Table 3. Significantly enriched terms (adjusted p-value < 0.01) after gene enrichment analysis for colorectal cancer cells. Enrichment done using the 50 genes most strongly correlated with the potential.**

### From 3D to 4D

| Source Type \ Predicted Type | Early_Neural | MN | Mesoderm | NMP | Neural | pMN |
|---|---|---|---|---|---|---|
| Early_Neural | 0.000 | 0.000 | 0.000 | 0.000 | 0.000 | 0.000 |
| FP | 0.000 | 0.000 | 0.000 | 0.000 | 0.000 | 0.000 |
| MN | 0.000 | 0.000 | 0.000 | 0.000 | 0.000 | 0.000 |
| Mesoderm | 0.000 | 0.000 | 0.000 | 0.000 | 0.000 | 0.000 |
| NMP | **0.035** | 0.000 | **0.331** | **0.140** | **0.487** | **0.007** |
| Neural | 0.000 | 0.000 | 0.000 | 0.000 | 0.000 | 0.000 |
| V3 | 0.000 | 0.000 | 0.000 | 0.000 | 0.000 | 0.000 |
| p3 | 0.000 | 0.000 | 0.000 | 0.000 | 0.000 | 0.000 |
| pMN | 0.000 | 0.000 | 0.000 | 0.000 | 0.000 | 0.000 |

### From 4D to 5D

| Source Type \ Predicted Type | Early_Neural | FP | MN | Mesoderm | Neural | V3 | pMN |
|---|---|---|---|---|---|---|---|
| Early_Neural | 0.000 | 0.000 | 0.000 | **0.001** | **0.001** | 0.000 | 0.000 |
| FP | 0.000 | 0.000 | 0.000 | 0.000 | 0.000 | 0.000 | 0.000 |
| MN | 0.000 | 0.000 | 0.000 | 0.000 | 0.000 | 0.000 | 0.000 |
| Mesoderm | 0.000 | 0.000 | 0.000 | **0.285** | **0.001** | 0.000 | **0.022** |
| NMP | 0.000 | 0.000 | 0.000 | 0.000 | 0.000 | 0.000 | 0.000 |
| Neural | 0.000 | 0.000 | **0.061** | **0.077** | **0.118** | 0.000 | **0.433** |
| V3 | 0.000 | 0.000 | 0.000 | 0.000 | 0.000 | 0.000 | 0.000 |
| p3 | 0.000 | 0.000 | 0.000 | 0.000 | 0.000 | 0.000 | 0.000 |
| pMN | 0.000 | 0.000 | 0.000 | 0.000 | 0.000 | 0.000 | 0.000 |

### From 5D to 6D

| Source Type \ Predicted Type | FP | MN | Mesoderm | V3 | p3 | pMN |
|---|---|---|---|---|---|---|
| Early_Neural | 0.000 | 0.000 | 0.000 | 0.000 | 0.000 | 0.000 |
| FP | 0.000 | 0.000 | 0.000 | 0.000 | 0.000 | 0.000 |
| MN | 0.000 | **0.013** | 0.000 | **0.001** | 0.000 | **0.006** |
| Mesoderm | 0.000 | 0.000 | **0.644** | 0.000 | 0.000 | **0.015** |
| NMP | 0.000 | 0.000 | 0.000 | 0.000 | 0.000 | 0.000 |
| Neural | 0.000 | 0.000 | 0.000 | 0.000 | 0.000 | 0.000 |
| V3 | 0.000 | 0.000 | 0.000 | 0.000 | 0.000 | 0.000 |
| p3 | 0.000 | 0.000 | 0.000 | 0.000 | 0.000 | 0.000 |
| pMN | **0.016** | **0.004** | 0.000 | 0.000 | **0.008** | **0.294** |

### From 6D to 7D

| Source Type \ Predicted Type | FP | MN | Mesoderm | V3 | p3 | pMN |
|---|---|---|---|---|---|---|
| Early_Neural | 0.000 | 0.000 | 0.000 | 0.000 | 0.000 | 0.000 |
| FP | **0.010** | 0.000 | 0.000 | 0.000 | 0.000 | 0.000 |
| MN | 0.000 | **0.008** | 0.000 | **0.001** | 0.000 | 0.000 |
| Mesoderm | 0.000 | 0.000 | **0.314** | 0.000 | 0.000 | **0.017** |
| NMP | 0.000 | 0.000 | 0.000 | 0.000 | 0.000 | 0.000 |
| Neural | 0.000 | 0.000 | 0.000 | 0.000 | 0.000 | 0.000 |
| V3 | 0.000 | 0.000 | 0.000 | 0.000 | 0.000 | 0.000 |
| p3 | 0.000 | 0.000 | 0.000 | **0.001** | **0.003** | 0.000 |
| pMN | **0.019** | **0.014** | 0.000 | **0.005** | **0.135** | **0.474** |

### From 7D to 8D

| Source Type \ Predicted Type | FP | MN | Mesoderm | V3 | p3 | pMN |
|---|---|---|---|---|---|---|
| Early_Neural | 0.000 | 0.000 | 0.000 | 0.000 | 0.000 | 0.000 |
| FP | **0.050** | 0.000 | 0.000 | 0.000 | **0.001** | 0.000 |
| MN | 0.000 | **0.008** | 0.000 | 0.000 | 0.000 | 0.000 |
| Mesoderm | 0.000 | 0.000 | **0.113** | **0.001** | **0.001** | **0.002** |
| NMP | 0.000 | 0.000 | 0.000 | 0.000 | 0.000 | 0.000 |
| Neural | 0.000 | 0.000 | 0.000 | 0.000 | 0.000 | 0.000 |
| V3 | 0.000 | 0.000 | 0.000 | **0.061** | 0.000 | 0.000 |
| p3 | **0.001** | 0.000 | 0.000 | **0.032** | **0.135** | **0.002** |
| pMN | **0.019** | **0.008** | 0.000 | **0.016** | **0.294** | **0.254** |

**Supplementary Table 4. Transition probabilities between cell types across consecutive time points in embryonic stem cells (Dataset 3). Probabilities are normalized globally per time-point transition.**

| FP Traj | | | MN Traj | | | V3 Traj | | |
|---|---|---|---|---|---|---|---|---|
| Rank | gene | scores | Rank | gene | scores | Rank | gene | scores |
| 1 | Ntn1 | 6.535e+01 | 1 | Srrm4 | 2.785e+01 | 1 | Elavl4 | 6.027e+01 |
| 2 | Gpc3 | 5.807e+01 | 2 | Chd3 | 2.728e+01 | 2 | Srrm4 | 5.933e+01 |
| 3 | Zfp36l1 | 5.695e+01 | 3 | Hecw2 | 2.657e+01 | 3 | Nfasc | 5.645e+01 |
| 4 | Col18a1 | 5.631e+01 | 4 | Robo2 | 2.624e+01 | 4 | Kalrn | 5.493e+01 |
| 5 | Fndc3b | 5.433e+01 | 5 | Elavl4 | 2.607e+01 | 5 | Elavl3 | 5.394e+01 |
| 6 | Hmga2 | 5.329e+01 | 6 | Trp53i11 | 2.526e+01 | 6 | Dpysl3 | 5.310e+01 |
| 7 | Tcf7l1 | 5.291e+01 | 7 | Nfasc | 2.497e+01 | 7 | Soga3 | 5.210e+01 |
| 8 | Pdzrn3 | 5.287e+01 | 8 | Nrp2 | 2.354e+01 | 8 | Dst | 5.147e+01 |
| 9 | Peg3 | 5.201e+01 | 9 | Elavl2 | 2.306e+01 | 9 | Myo16 | 5.134e+01 |
| 10 | Tcf7l2 | 5.173e+01 | 10 | Isl1 | 2.264e+01 | 10 | Stxbp1 | 5.110e+01 |
| 11 | Rfx4 | 4.985e+01 | 11 | Gng2 | 2.257e+01 | 11 | Tubb3 | 5.017e+01 |
| 12 | Hells | 4.865e+01 | 12 | Myo16 | 2.201e+01 | 12 | Trp53i11 | 4.910e+01 |
| 13 | Rif1 | 4.859e+01 | 13 | Stk32a | 2.186e+01 | 13 | Elavl2 | 4.902e+01 |
| 14 | Anp32b | 4.843e+01 | 14 | Cntn2 | 2.173e+01 | 14 | Rbfox3 | 4.876e+01 |
| 15 | Sox6 | 4.822e+01 | 15 | Rtn4 | 2.158e+01 | 15 | Onecut2 | 4.814e+01 |
| 16 | Dek | 4.789e+01 | 16 | Dpysl3 | 2.140e+01 | 16 | Ebf3 | 4.813e+01 |
| 17 | Nasp | 4.772e+01 | 17 | Dst | 2.136e+01 | 17 | Myo1b | 4.790e+01 |
| 18 | Nxn | 4.765e+01 | 18 | Cracd | 2.136e+01 | 18 | Cpeb4 | 4.780e+01 |
| 19 | Pola1 | 4.736e+01 | 19 | Onecut1 | 2.128e+01 | 19 | Srrm3 | 4.771e+01 |
| 20 | Plekha7 | 4.735e+01 | 20 | Ina | 2.078e+01 | 20 | Scg3 | 4.767e+01 |
| 21 | Notch2 | 4.669e+01 | 21 | Elavl3 | 2.012e+01 | 21 | Clvs1 | 4.738e+01 |
| 22 | Itgb8 | 4.614e+01 | 22 | Tspan5 | 2.010e+01 | 22 | Malat1 | 4.660e+01 |
| 23 | Ccnd2 | 4.596e+01 | 23 | Slit3 | 2.008e+01 | 23 | Map1a | 4.615e+01 |
| 24 | Wls | 4.560e+01 | 24 | Rbfox3 | 2.007e+01 | 24 | Crmp1 | 4.610e+01 |
| 25 | Cux1 | 4.559e+01 | 25 | Cers6 | 1.997e+01 | 25 | Stmn2 | 4.604e+01 |

**Supplementary Table 5. Top 25 differentially expressed genes in each trajectories (FP, MN, and V3) in embryonic stem cells (Dataset 3), ranked by decreasing Wilcoxon rank-sum test statistic.**
